# Ecosystem size-induced environmental fluctuations affect the temporal dynamics of community assembly mechanisms

**DOI:** 10.1101/2022.01.10.475621

**Authors:** Raven L. Bier, Máté Vass, Anna J. Székely, Silke Langenheder

## Abstract

Understanding processes that determine community membership and abundance is important for many fields from theoretical community ecology to conservation. However, spatial community studies are often conducted only at a single timepoint despite the known influence of temporal variability on community assembly processes. Here we used a spatiotemporal study to determine how environmental fluctuation differences induced by mesocosm volumes (larger volumes were more stable) influence assembly processes of aquatic bacterial metacommunities along a press disturbance gradient. By combining path analysis and network approaches, we found mesocosm size categories had distinct relative influences of assembly process and environmental factors that determined spatiotemporal bacterial community composition, including dispersal and species sorting by conductivity. These processes depended on, but were not affected proportionately by, mesocosm size. Low fluctuation, large mesocosms primarily developed through the interplay of species sorting that became more important over time and transient priority effects as evidenced by more time-delayed associations. High fluctuation, small mesocosms had regular disruptions to species sorting and greater importance of ecological drift and dispersal limitation indicated by lower richness and higher taxa replacement. Together, these results emphasize that environmental fluctuations influence ecosystems over time and its impacts are modified by biotic properties intrinsic to ecosystem size.

## Introduction

The community composition of both micro- and macro-organisms at a given point in space and time results from the interaction of multiple assembly processes, including ecological drift, species sorting (environmental filtering), dispersal, and speciation (1-5). Most observational metacommunity studies, however, focus only on spatial snapshots without considering temporal dynamics of community assembly and association networks, or historical contingencies (2, 6). Hence, we still lack knowledge about the underlying mechanisms and regulating factors that temporal dynamics encompass.

When species sorting assembles communities, their composition tracks changes in environmental conditions that occur in time and space (2, 7). However, environmental tracking can be hindered or disrupted (8). Such asynchrony can lead to historical contingencies by priority effects (e.g., (8, 9), which can occur during early community formation or when communities re-assemble following perturbation. An important consequence of priority effects is that they impede or delay environmental tracking enacted by species sorting.

Environmental changes may influence temporal community assembly processes and the strength of this can be regulated by ecosystem size (e.g., 6). Studies have shown that microbial communities exposed to disturbances are initially, and often to a strong degree, stochastically assembled, but that the importance of species sorting increases later during community re-assembly as more species from the regional species pool arrive (11-14). Rapidly fluctuating environmental conditions, however, may continuously disrupt environmental tracking by reducing opportunities for species sorting to select and shape local communities before the environmental conditions change again. This might promote coexistence of species with different niche optima (20-22) and, thus, reduce beta diversity (16), or could cause extinctions that bolster dispersal limitation and priority effects (23). Nevertheless, many studies happen in controlled settings; thus, we lack knowledge on the temporal dynamics of these processes within larger, more complex habitats which track environmental changes (2, 17). Disturbance strength may uniquely affect microbial communities in ecosystems of different sizes as ecosystem size may influence assembly processes by increasing habitat heterogeneity, community abundance (6, 18, 19), and the pace at which communities track environmental changes. For instance, communities may experience different environmental variability including press disturbances (e.g., climate warming, eutrophication, or saltwater incursion), periodic and stochastic environmental fluctuations, where the latter may influence community assembly in response to the former over time and space.

Here, we implemented an experiment with freshwater bacterial metacommunities to test how different ecosystem size-induced environmental fluctuations influence the temporal dynamics of community assembly mechanisms. We collected a 64-day time series from mesocosms that allowed bacterial communities sufficient time to experience natural environmental fluctuations. Specifically, we set-up a natural experimental landscape with mesocosms containing identical lake water that differed in volume, which induced differences in environmental fluctuation intensity among the mesocosms. We created a press disturbance by applying a salinity gradient to each set of mesocosm volumes as it has been shown that salinity affects bacterial communities in many ecosystems (e.g., (10-13). We hypothesized that the importance of species sorting would increase over time in local communities of larger mesocosms that experience relatively minor environmental fluctuations because their communities will have sufficient time for species selection in response to the initial salinity. Second, other environmental changes occurring in mesocosms would be slow in large mesocosms and this would allow time for taxa to be recruited from internal and external dispersal sources and to become active. We expected that species sorting related to salinity differences across communities, i.e., at the metacommunity scale, promotes recruitment of taxa best suited to the salinity. Last, we hypothesized that stochastic and/or dispersal-related assembly processes should be more important in small mesocosms where communities experience strong environmental fluctuations that continually disrupt environmental tracking. We combined quantitative path analysis methods that aim to estimate metacommunity processes with a network approach that identifies environmental tracking patterns through local and time-delayed co-occurrences to provide insights into temporal dynamics of microbial ecosystems (14).

## Methods

### Experimental set-up

Three different sizes (24.5, 70, or 200 L) of hard-shell polyethylene mesocosms were arranged in a field beside Lake Erken (16 per size category) and filled with 0.1 mm filtered lake water from Lake Erken in Sweden (59°51’N 18°35’E) (water properties in Supplementary Information). Mesocosms were seeded with 1 L of sieved and mixed surface sediments collected from Lake Erken at ∼0.5 m water depth.

To induce species sorting with a press disturbance, a salinity gradient was created within each mesocosm size category using nitrate- and phosphate-free sea salt (Red Sea Aquatics Ltd, Verneuil-sur-Avre, France) and ranged from freshwater (0 ‰) to 6 ‰ with 0.4 ‰ increments (rationale for range in Supplementary Information). Mesocosm water surface area and volume were proportional such that air or rain dispersal was proportional across size classes. Mesocosm sediment was also a recruitment source (15-17). Equal mesocosm bottom surface areas allowed for equal recruitment independent of fluctuation category.

### Monitoring and sampling

Mesocosms were monitored on days 1, 2, and 4, and then every fourth day for 64 days from July to September 2016. Monitoring included depth profiles of conductivity (to measure salinity changes) and temperature, and depth integrated pH, chlorophyll-*a*, and colored dissolved organic matter (CDOM) fluorescence (see Supplementary Information for details). Weather data from Svanberga, Sweden (0.87 km southwest of the site) included daily precipitation and hourly air temperature (Swedish Meteorological and Hydrological Institute). Every eighth day, water was collected for total organic carbon (TOC), total nitrogen (TN), and total phosphorus (TP) and analyzed using established methods (18).

Water samples for enumerating microorganism cells were collected simultaneously with bacterial community composition (below) and preserved with sterile formaldehyde to 2.5 % (19). Samples were stained with SYTO™ 13 Green Fluorescent Nucleic Acid Stain (ThermoFisher Scientific), counted (CyFlow Space flow cytometer, Partec, Münster, Germany) and analyzed using FlowingSoft software (Perttu Terho). Total community size was calculated as cell abundance (mL^-1^) multiplied by mesocosm volume.

### Bacterial community composition

Mesocosm water was collected on days 1, 2, 4, 8 and every 8^th^ day thereafter for 64 days to assess community composition through 16S rRNA amplicon sequencing to specifically detect active members (20). Depth integrated 0.5 L water samples, and air and rain immigration samples (see Supplementary Information), were collected and filtered onto 0.2 μm pore-size filters (47 mm Supor-200 filters, Pall Corporation, Hampshire, UK) until 5 minutes or 0.5 L volume was reached. Filters were flash-frozen in liquid nitrogen and stored at −80 ºC. DNA from initial lake water and sediment used in the experiment were sampled to learn initial communities and seed banks.

Nucleic acids were extracted using a modified protocol from Easy-DNA™ kit (Invitrogen, Carlsbad, CA, USA). See Supplementary Information and DOI for a detailed protocol (dx.doi.org/10.17504/protocols.io.xekfjcw). Samples were submitted to SNP&SEQ Technology platform at SciLife in Uppsala, Sweden for two Illumina MiSeq PE300bp sequencing runs with v3 chemistry.

### Data processing

Sequencing resulted in 35.6 million paired reads from 609 demultiplexed samples including 12 extraction and PCR negatives. Primers were removed from sequences using cutadapt v 2.7 ref. (21). The DADA2 pipeline (22) was used for sequence processing and taxonomy assignment of Amplicon Sequence Variants (ASVs) using the SILVA v. 138.1 reference database (23) (Supplementary Information, Table S1).

For beta diversity analyses, ASVs with counts less than 10 were removed and samples were subsampled to a minimum of 5 028 reads, retaining 7 983 unique ASVs. Samples not meeting the 5 028 reads requirement were excluded (Table S2). Both alpha and beta diversity datasets represented 99 % coverage. Raw sequences are available in the European Nucleotide Archive (study accession number PRJEB26595).

### Statistical analyses

Statistical analyses were conducted in R (v3.4.3 and v4.0.2) ref. (24) with package “vegan” (25) unless otherwise specified.

#### Fluctuation magnitudes among mesocosm sizes

For each environmental variable, fluctuations data were analyzed using the mean of absolute differences of mesocosms in a size category between one date and the previous sampling date. For variables with depth profiles (conductivity and temperature), the absolute difference at each depth was used to calculate the mean change per mesocosm. The environmental variables dataset is in the DiVA repository (26). To determine if the magnitude of changes differed between mesocosm sizes over time, nonparametric tests for repeated measures with an ANOVA-type statistic (ATS) were used (R package and function *nparLD*, ref. (27)). Mesocosms were assessed using principal components analysis (PCA) of original and absolute changes of environmental variables (both log-transformed) and fit with environmental vectors (Fig. S2).

#### Community composition, diversity, and recruitment

Non-metric multidimensional scaling (NMDS) with Bray-Curtis dissimilarities was used to visualize bacterial community composition and environmental variables. Shannon’s index and Pielou’s evenness were calculated and richness was estimated using the package “breakaway” (28). Temporal beta diversity differences in each mesocosm were evaluated by comparing each community with the previous using Jaccard pairwise dissimilarity values. Dissimilarity was partitioned between taxa turnover (taxa replacement) and community nestedness (chronological subsets of taxa) using package “betapart” v1.5.2 ref. (29). Variation from each partition captured by mesocosm size was compared using PERMANOVA tests (30) with function *adonis* and 999 permutations.

Recruitment was evaluated by pooling each mesocosm’s active ASVs across days; ASVs present on day one were removed from the pool leaving those recruited during the experiment. Recruited ASVs were matched with their source seed bank(s) based on DNA from sediment, initial lake water, air, and rain. Sources for unmatched ASVs were considered unknown. For each mesocosm, the percent of recruited ASVs was calculated, split into each source, and examined across the salinity gradient using Pearson’s correlations.

#### Path analysis

To detect drivers of metacommunity dynamics, a spatiotemporal path analysis was used (31). This method calculates dissimilarity for all community pairs sampled over time and space and estimates, as individual paths on this beta-diversity measure, the influences of spatial distance (Δx), temporal distance (Δt), environmental distance (ΔE), mean community size (<J>, cell abundance multiplied by mesocosm volume), and absolute differences in community size (ΔJ) and taxa richness (ΔS). Nestedness between sites should explain a positive link between differences in community size and richness thereby increasing community dissimilarity (31). Bray-Curtis dissimilarity was used for the community dissimilarity matrix (β_bc_). A permutation-based approach adjusted with Benjamini-Hochberg procedure indicated path significance. Model fit was assessed with the standardized root mean square residual (SRMR). The analysis was run separately for each mesocosm size using the days required for network analysis (Supplementary Information), with the *sem* function in R package “lavaan” (32).

#### Network analysis

To uncover local and time-delayed microbial associations and the extrinsic effects of environmental variables on bacteria, extended local similarity analysis (eLSA) was applied (14). Given our temporal data, this approach detects undirected associations (e.g., without time delays), and associations where the change of one factor (a taxon or environmental variable) chronologically leads or follows another factor. For a link between taxa and environmental variables, the association type (delayed or non-delayed) can indicate tracking that is time-lagged due to transient priority effects, or simultaneous through species sorting, respectively. Associations were determined for each mesocosm size using eLSA wherein mesocosms from each size category were used as fluctuation level replicates (n = 16). Because of the within- and across-size variability of bacterial communities (e.g., significant differences in taxa richness), we selected and analyzed only the core bacterial groups for each mesocosm size to make it comparable. Hence, networks used the 50 most abundant ASVs from each size category. eLSA (v1.0.2) was run over eight sampling points, allowing for local similarity (LS) correlations between samples taken eight days apart (d = 1). LS correlations (LS value ≥ 0.05; Q ≤ 0.01) were visualized in Cytoscape v3.8.2 (33). Network characteristics were calculated using the Cytoscape plugin NetworkAnalyzer (34). See Supplementary Information for details on sample selection, dominant ASV abundances, and statistics.

## Results

### Environmental fluctuations in mesocosms

Environmental variable fluctuations corresponded with mesocosm size and reflected rainfall and air temperature (Fig. S1, Table S3, Fig. S2). Size categories experienced significantly different conductivity and temperature fluctuations. After four days small and medium mesocosm conductivity fluctuated more than large mesocosms (Fig. S1, Table S3). Mean temperature fluctuation increased inversely with mesocosm size (Table S3). Mesocosm depth profiles showed stable conductivity, but temperature decreased with depth in medium and large mesocosms (Fig. S3).

Mesocosm sizes differed in nutrient concentrations and the absolute change of other environmental variables (chl-*a*, CDOM, pH, TN, TOC, TP and cell abundance, Table S3) and most pairwise comparisons showed that the degree of change differed significantly between sizes with the greatest changes in small mesocosms. Absolute changes between sampling dates and individual timepoints grouped according to size (Fig. S2). Measured nutrients and conductivity positively correlated with decreasing mesocosm sizes (environmental vector correlations, *p* < 0.05). For water temperature, sampling date was more influential than mesocosm size. Cell abundances (cells mL^-1^) increased over time and was highest in small and medium mesocosms (ATS, *p* < 0.001, Fig. S4A). From day 24, total community abundance per mesocosm was lower in small than medium and large mesocosms (ATS, all *p* < 0.002, Fig. S4B).

### Community composition, diversity, and recruitment

Bacterial community composition shifted with time and initial conductivity in all mesocosm sizes (Fig. S5). Diversity indices (estimated richness, Pielou’s evenness, Shannon’s index) did not differ on the first day (two-way ANOVA, all *p* > 0.05), but over time all three indices differed by mesocosm size (Fig. 1, repeated measures ANOVA, all overall *p* ≤ 0.001; pairwise Bonferroni adjusted). All sizes differed significantly in bacterial richness which was lowest in small and highest in large mesocosms (all *p* ≤ 0.001). The more evenly distributed ASV abundances in large mesocosms widened the separation in Shannon’s index diversity between large and small or medium mesocosms, indicating a greater presence of dominant and/or rare taxa in smaller mesocosms (Fig. 1). Mesocosm size explained some variability in beta diversity from turnover (F model = 13, R^2^ = 0.04, *p* ≤ 0.001) with communities in small mesocosms experiencing higher turnover by taxa replacement than those in large mesocosms (Wilcoxon Test, W = 4276, Bonferroni *p*.*adj*. = 0.02, Fig. S6A). Statistically, mesocosm size did not explain variability in nestedness, although communities in large mesocosms trended towards greater nested species loss (Fig. S6B).

**Figure 1.**
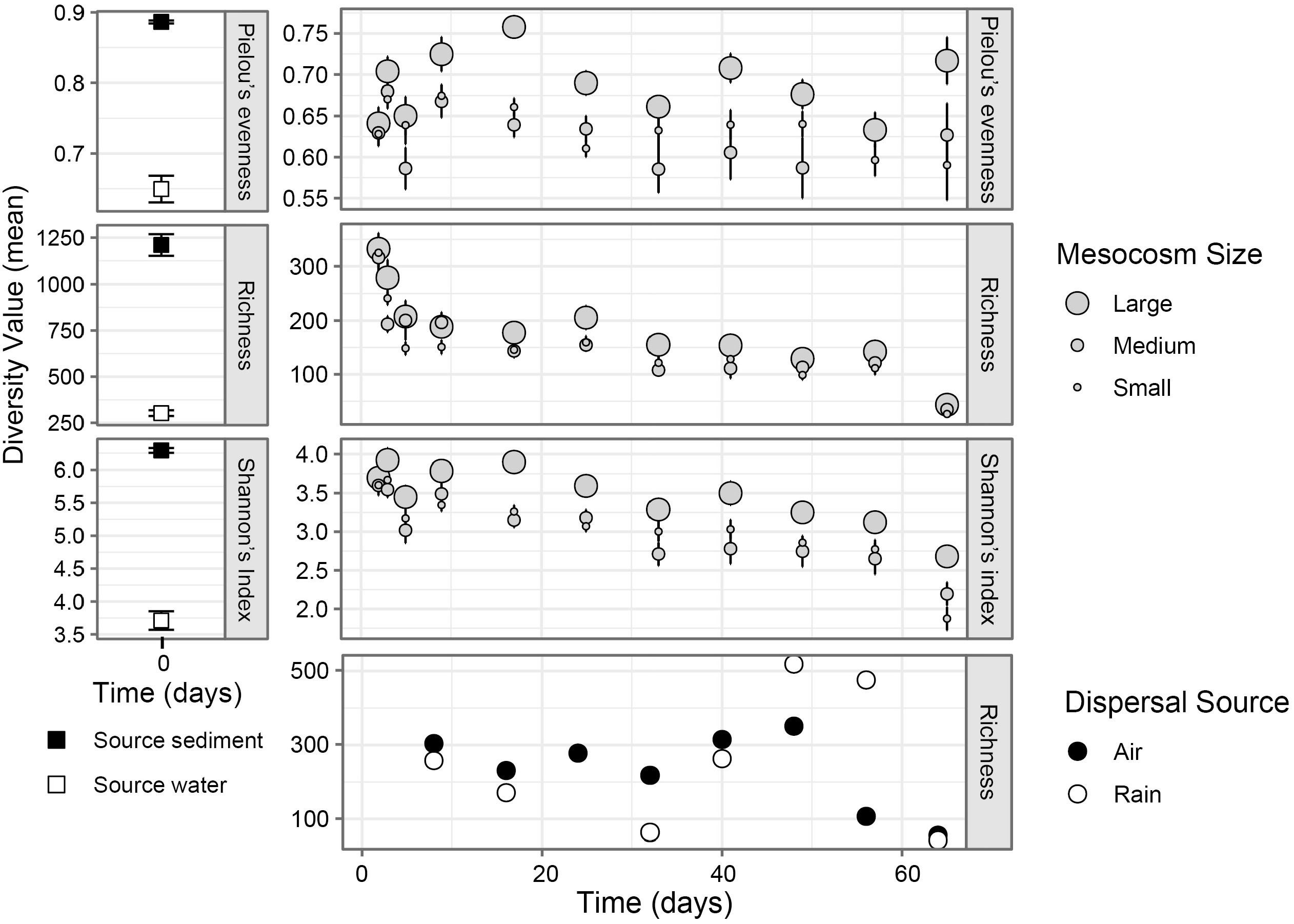
Temporal patterns of alpha diversity metrics for bacterial communities in dispersal sources (air and rain), source media (sediment and water) (DNA) and mesocosm water (RNA). Error bars are standard error. Diversity metrics for large mesocosms greater than both small and medium mesocosms (repeated measures ANOVA, pairwise t-test with Bonferroni correction, p < 0.05). Source media n = 3, mesocosm sizes n = 16, air and rain n = 1. Note difference in y-axis scales.

Recruited ASVs as a percentage of total unique ASVs, had weak negative correlations with the salinity press disturbance in small and large mesocosms (r = -0.35 and -0.39, *p* < 0.005, respectively, Fig. S7). Less than 15 % of recruited ASVs in each mesocosm were attributed to a known source. In all mesocosm sizes, recruitment from water declined significantly with initial salinity (small: r = -0.90, medium: r = -0.85, large: r = -0.89, all *p* < 0.001). Recruitment from sediment showed different patterns across salinity levels in small and large mesocosms: it decreased in small mesocosms and was unchanged in large mesocosms (r = -0.70, *p* = 0.002; r = 0.45, *p* = 0.08, respectively). Sediment was typically the largest recruitment source in the most saline mesocosms. Air and rain recruitment was related to salinity level only in the medium mesocosms where it was weakly positively correlated (air: r = 0.52, *p* = 0.04, rain: r = 0.58, *p* = 0.02).

### Path analysis

Bacterial metacommunities in mesocosms of different sizes experienced disparate relative influences from species sorting by environmental variation, demographic stochasticity, and dispersal limitation (Fig. 2). The model fit for small mesocosms was roughly twice that of medium and large mesocosms (Fig. 2).

**Figure 2.**
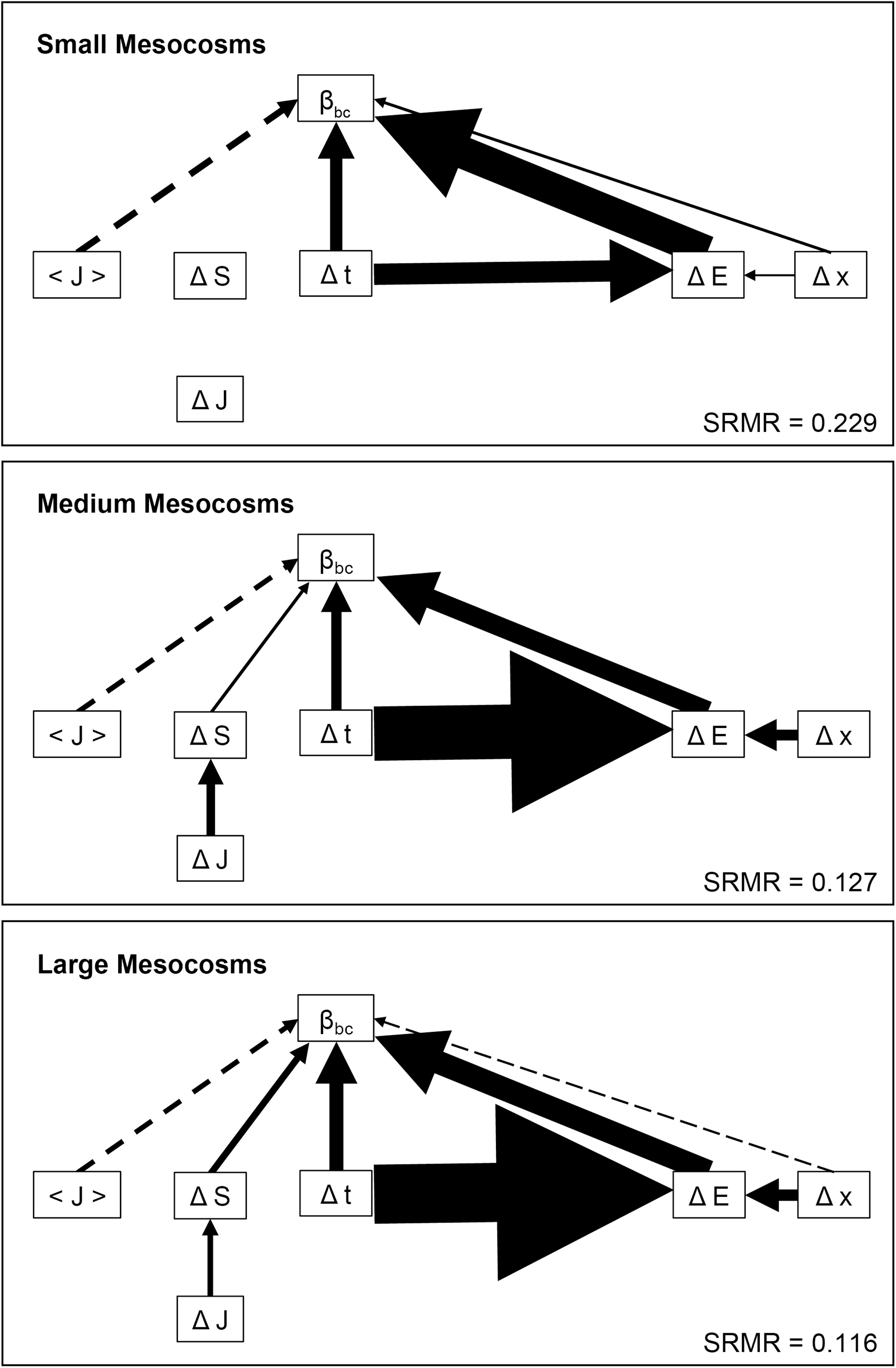
Path analysis diagrams of small, medium, and large mesocosm sizes. The influence of spatial distances (Δx), temporal distances (Δt), environmental distances (ΔE), mean community size (<J>), absolute difference in community size (ΔJ) and species richness (ΔS) on community dissimilarity (β_bc)_ was quantified following Jabot et al.’s framework (2020). Arrow width represents standardized estimate strength with positive estimate arrows in solid lines and negative estimates in dashed lines. For environmental variables, the absolute values of standardized estimates were added. Effects shown have p < 0.05. SRMR = Standardized Root Mean Square Residual. See Tables S4-S6 for standardized estimate values.

Species sorting (ΔE) had the most influential direct effect on community dissimilarity (β_bc_) (Fig. 2). This effect was strongest in small mesocosms and similar in medium and large mesocosms (sum of absolute standardized estimates 0.925, 0.773, and 0.766, respectively), but all sizes had significant environmental distance and community dissimilarity relationships (Tables S4-S6). Small mesocosms had five significant relationships between community dissimilarity and environmental variables (conductivity, temperature, chlorophyll-*a*, TOC, and TN); large mesocosms had three (conductivity, temperature, and chlorophyll-*a*), and medium mesocosms had only conductivity. Conductivity correlated most strongly with community dissimilarity of medium, followed by large and small mesocosms. Significant correlations between temporal (Δt) or spatial (Δx) distance and environmental (ΔE) distance were positive and increased with mesocosm size. The indirect effect of time on community dissimilarity through species sorting was apparent with all measured variables except conductivity.

Demographic stochasticity was indicated by significant negative relationships between mean community size (<J>) and community dissimilarity in all mesocosm sizes (Fig. 2, Tables S4-S6). Small mesocosms had the strongest influence by demographic stochasticity. All mesocosm sizes had positive correlations between temporal distance and community dissimilarity indicating additional demographic stochasticity. Relationship strengths differed with size: temporal changes had the greatest influence in large, then small, then medium mesocosms.

Dispersal limitation shown as a positive correlation between geographic distance and community dissimilarity appeared only for small mesocosms (Fig. 2). Large mesocosms had a significant negative correlation between geographic distance and community dissimilarity but this was considered an artefact of the linear modelling framework (31) and negligible compared with the relationship between space and community dissimilarity via the environmental variation pathway.

The path analysis for medium and large mesocosms also suggested an effect of taxa nestedness whereby communities form as subsets of original communities over time or space (Tables S4-S6). First, differences in community richness (ΔS) positively correlated with community dissimilarity. This relationship was strongest in large mesocosms. Second, a significant positive relationship occurred between differences in community size (ΔJ) and richness in medium and large mesocosms.

### Association networks

Association networks of the 50 most abundant ASVs (members of Actinobacteriota, Bacteroidota, Cyanobacteria, Planctomycetota and Proteobacteria) differed among the three mesocosm sizes (Fig. 3, Table S7). The number of total edges and ASV nodes increased with mesocosm size, and the proportion of delayed (time-shifted) associations were higher in larger mesocosms (small: 25.9 %, medium: 49.8 %, large: 46 %) (Table S7). Small mesocosms had the most ASVs (n = 18) that were unassociated with environmental variables and bacterial abundance while medium and large mesocosms had only 4 and 6 ASVs, respectively (Table S7). Network subsets showed no connection between conductivity and ASVs of small mesocosms, while conductivity influenced many abundant ASVs from medium and large mesocosms (mainly phylum Proteobacteria). In large mesocosms, conductivity had a direct (non-delayed) influence on ASVs (except one Cyanobacterium), while in medium mesocosms it had both time-shifted (e.g., mainly positive in Bacteroidota and negative in Proteobacteria) and non-delayed (e.g., Cyanobacteria) associations with taxa (Fig. S8, Table S8).

**Figure 3.**
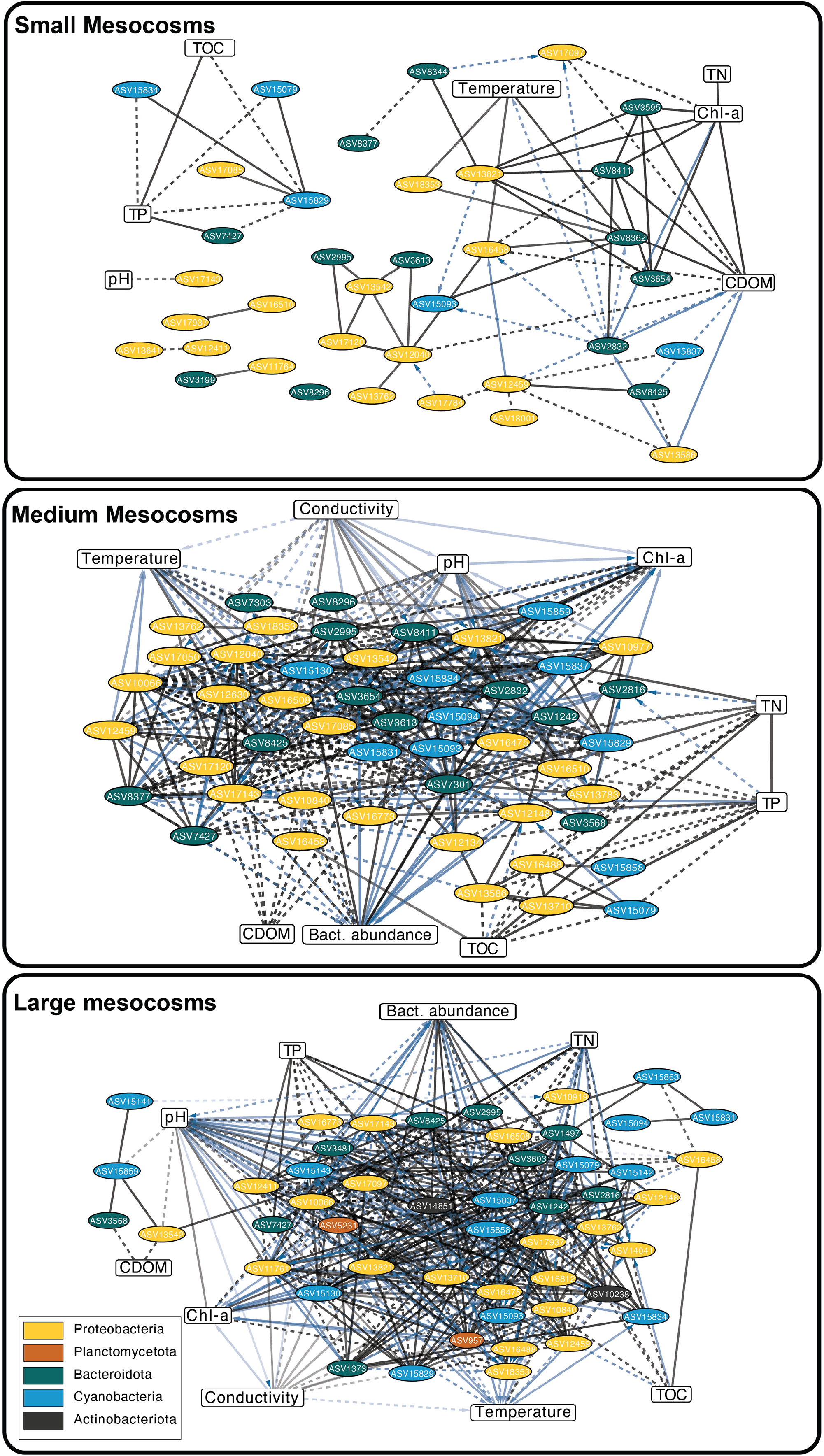
Association networks and the relative abundances of the 50 most abundant bacteria of the three mesocosm size categories (n = 16). All significant ((p ≤ 0.01 and Q ≤ 0.01) pairwise local similarity (LS) correlations ≥ 0.05 are shown as edges in the networks. Each node represents an ASV (ellipse) or an environmental factor (rectangle). Edge transparency is proportional to the association strength (based on LS values). Solid lines refer to positive associations while dashed lines to negative ones. Edge colors indicate delayed (blue) and non-delayed (black) associations between ASVs and/or environmental variables. Arrows point toward the lagging node.

Association networks were quantitatively compared by mesocosm size with commonly used topological characteristics. Negative associations, average number of neighbors, and network density (the proportion of possible edges that are associated with nodes) increased with mesocosm size (Table S7). Further, small and medium mesocosms networks were less centralized (the concentration of centrality among the nodes) than those in large mesocosms. When considering only taxa associations, small mesocosms had the least centralized network with more taxa displaying similar numbers of links (Table S7).

## Discussion

Here we show how differences in environmental fluctuation strengths due to differences in ecosystem, i.e. mesocosm, size influenced the temporal dynamics of community assembly in response to a salinity press disturbance (Fig. 4). First, species sorting was generally the most influential process for all mesocosms but differences in how species sorting operated among mesocosm sizes at the community (path analysis) and individual taxa levels (association network analysis). These evaluations indicated that under low environmental fluctuations, dominant ASV populations were effective trackers of environmental conditions. When ecosystem size-induced environmental fluctuations were strong (i.e., small mesocosms), environmental tracking was disrupted. Second, the salinity press disturbance initiated environmental tracking, especially under stable conditions (i.e., larger mesocosms), through the recruitment of taxa from seed banks (mainly sediment at high salinity). Third, stochasticity and dispersal-related assembly processes (e.g., dispersal limitation) generally were more important for communities of small ecosystems. Overall, our study aligns with previous findings that ecosystem size influences community assembly processes (35-37), but we identifed this effect to derive from environmental fluctuations created by ecosystem size differences and corresponding differences in species sorting effects.

**Figure 4.**
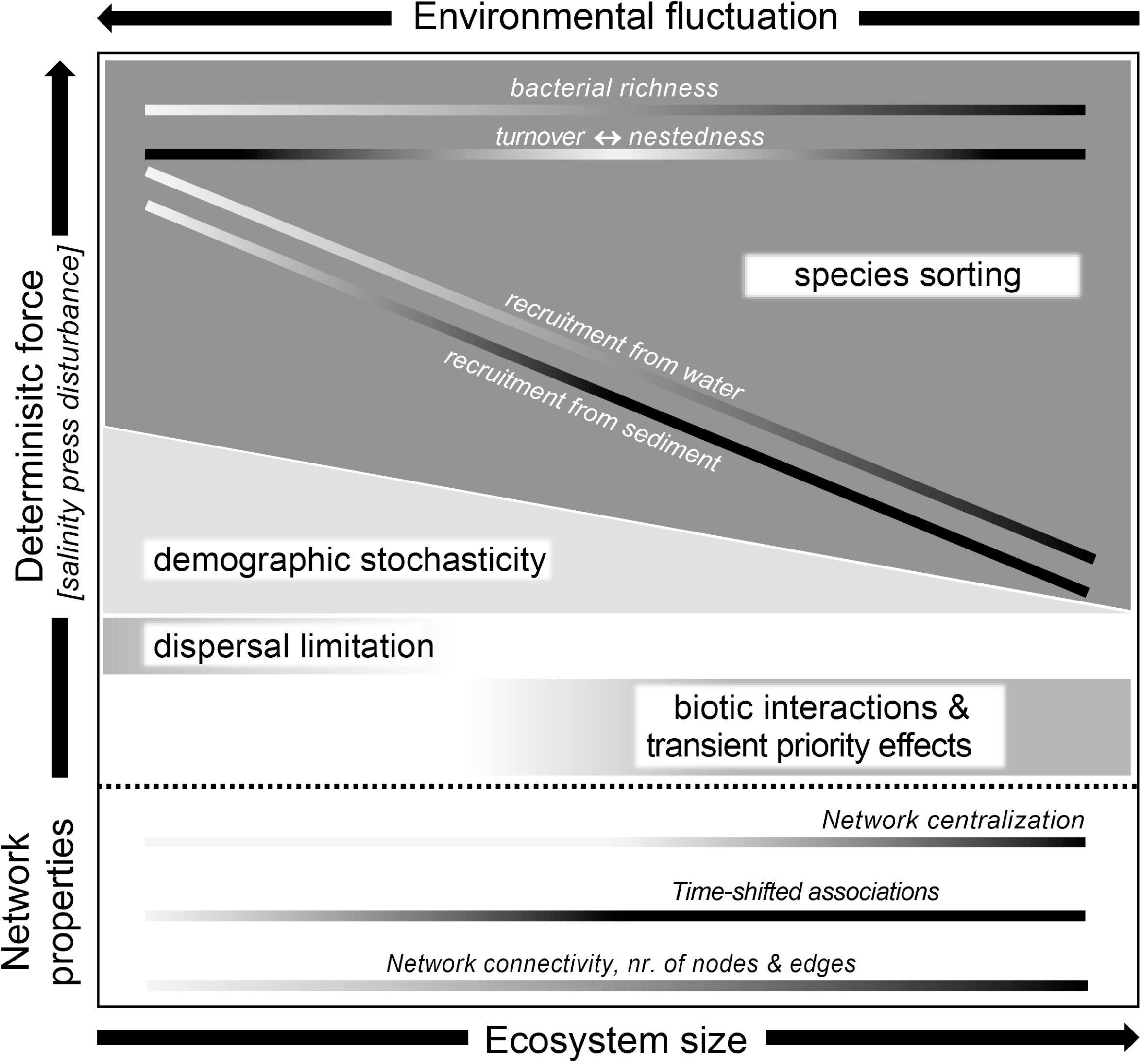
Conceptual figure for the interpretation of statistical results and patterns based on path analysis, network analysis, and the partitioning of beta-diversity. In our study, the dominant deterministic force was the applied salinity press disturbance. Ecosystem size was manipulated by different volumes of mesocosms. Darker shading in bars indicates greater influence of the process.

### Salinity press disturbance enforces environmental tracking

Differences in the magnitude of salinity press disturbances induced clear compositional shifts within and across mesocosms over time. This was expected as we used salinity to induce species sorting because it is an environmental factor that causes clear taxonomic differences in aquatic bacterial communities (13, 38-41). However, there were disparities in how well communities in each mesocosm size tracked temporal changes in salinity.

The path analysis and network analysis results indicated that species sorting patterns differed across mesocosm sizes and were altered by time. The direct effects of significant environmental variables with unidirectional influences (i.e., salinity and temperature) on species sorting were most influential in medium and large, stabler mesocosms. However, when variables prone to feedbacks (i.e., nutrients, see below) were included into the total environmental effect on composition, species sorting was greatest in small, highly fluctuating mesocosms. In contrast, the indirect effect of time on composition via species sorting increased with mesocosm size and was driven primarily by changes in all environmental variables except for salinity (which changed minimally within a mesocosm compared to the salinity gradient). This temporal pattern generally agreed with the network results of the 50 most abundant bacteria which showed that they best tracked multiple environmental variables over time in large and medium mesocosms. Here, almost all ASVs directly linked to environmental variables and populations of core groups of taxa oscillated correspondingly with temporal salinity changes. In addition, when we isolated taxa and salinity associations, the network approach revealed many non-delayed associations in large mesocosms, indicating that the most abundant bacterial populations rapidly (simultaneously) tracked changes. Despite the path analysis results, no associations were found in the small mesocosms which could otherwise indicate salinity tracking through time.

Several reasons may explain the contradiction between the two analytical approaches regarding direct species sorting. First, in the network analysis, we calculated associations only among the 50 most abundant taxa, thus, we likely overlook conditionally rare taxa that can be temporarily abundant (42) as a consequence of the rapidly changing environment in small mesocosms. This is supported by the trend of higher taxa turnover and direct demographic stochasticity (discussed below) in small mesocosms. Another explanation may include the phenomenon that bacterial communities can be an imprint of past environmental conditions (43) and the correlations detected between community dissimilarity and environmental variables might coincide with prior processes. Last, the analytical approaches generally agreed concerning direct effects by salinity, but differed for variables with the potential for feedbacks (i.e., nutrients). Although the path analysis portrays nutrients as effect variables, they are also modified by microorganisms. Likely due to the salinity and greater proportions of sediment, small mesocosms had greater nutrient concentrations and higher cell densities including from observed algal blooms. These conditions could increase competition which hinders synchrony between abiotic variables and taxa abundances (44).

Taken together, our findings (conceptualized in Fig. 4) around the importance of species sorting and the strong temporal influences highlight the distinct differences in the mechanisms underlying species sorting in mesocosms of different sizes. These findings became apparent through combining the path analysis, which captures both spatial and temporal patterns at the whole community level, and the network analysis, which captures time-associated patterns of the most abundant populations.

### Ecosystem size regulates community assembly and associations among bacterioplankton

While the different environmental conditions from the press disturbance and ongoing environmental changes throughout the experiment might explain why species sorting was the main driver of metacommunity assembly, our study suggests that other factors related to ecosystem size (e.g., spatial environmental heterogeneity) could further regulate metacommunities.

The importance of species sorting can increase with environmental heterogeneity, i.e., the number of niches that are available for colonization across patches (45, 46). In our study, large mesocosms contained greater spatial environmental heterogeneity evidenced by depth associated changes in temperature and light. Hence, across mesocosms spatial environmental heterogeneity could explain why species sorting was more apparent in large compared to small mesocosms based on the network analysis. This might also explain the increased associations in larger mesocosms, enhancing the probabilities for true biotic interactions. This increase may be attributed to (i) the greater availability of niches (and consistency of nutrients) found in larger mesocosms, or (ii) the synchronous establishment of bacteria which might have a better chance in a stable environment. In a study of protists experiencing light-dark fluctuations in aquatic microcosms and models, high fluctuations disrupted species synchrony between patches (47). Spatial heterogeneity could also explain the greater bacterial richness as ecosystem size increased.

Network topological features were partially influenced by mesocosm size: bacteria were more connected in medium or large than small mesocosms, suggesting that abundance dynamics were less similar across small mesocosms and indicating asynchrony among dominant bacteria. In these less densely populated, large mesocosms, competition may have lessened which can lead to greater synchrony between species that is driven by changes in abiotic conditions (44). In our mesocosms, more connected, centralized communities with greater network density occurred as size increased, indicating the presence of subnetworks or cliques and that, due to lower fluctuation strength, the establishment of more connected, denser networks were common under stabler environments (Fig. 4). Because size replicates in the network analysis spanned the salinity gradient, this further suggests that the spatial environmental heterogeneity of salinity had less importance for potential biotic associations among stabler mesocosms.

Taken together, we suggest that these patterns indicate mesocosm size-specific mechanisms of species sorting: in small mesocosms, changes in community composition from species sorting primarily occurred through taxa replacement in response to variation in multiple environmental factors. In contrast, in larger mesocosms, environmental change was more gradual and cascaded into compositional differences through abundant bacteria tracking environmental changes over time by changing in relative population size, with lower replacement (Fig. 4).

### Recruitment of the members of bacterioplankton

Initial community size due to differences in mesocosm volume might have affected the resulting community composition through species sorting, but other factors related to the experimental set-up were unlikely to have substantial influence. The experimental set-up ensured no extensive differences in the recruitment of novel species from external sources due to proportional mesocosm surface areas and equal initial sediment volumes. There were also no differences in estimated richness of active bacteria between mesocosms on the first day of the experiment. The dispersal sources (rain and air deposition as well as seed banks in sediments and lake water) all harbored high diversity and in previous studies were shown to be important recruitment sources for novel taxa following salinity disturbances (15-17) and other types of environmental change (48). Although large mesocosms contained more total microorganisms and thereby possibly a larger planktonic seed bank from which taxa could respond to the salinity disturbance, recruitment from the water seed bank declined with salinity in all mesocosm sizes (Fig. 4). Even with reduced dispersal, future studies that extend temporal sampling beyond the 64 days sampled here may eventually see eco-evolutionary processes such as increased tracking of environmental conditions in small mesocosms due to bacterial diversification, which can be intensified by a history of environmental adversity (49). The high percentage of ASVs for which we did not identify a source (85%) could indicate dispersal from other sources such as the snails we observed on most mesocosms. or the effect of sequencing depth which can miss the rarest taxa.

### Roles of stochasticity and dispersal-related processes

Demographic stochasticity (leading to ecological drift) was an important driver of community assembly of all mesocosms via community size with the strongest direct effect in small mesocosms (Fig. 4). This result is bolstered by previous studies showing that ecological drift more often occurs in small communities (50, 51) especially when the importance of species sorting is weak (52) or when the effective community size is small due to dispersal limitation (53). This may be why we did not detect synchronous environmental tracking from the dominant populations across the small mesocosms. Drift can also alter the outcome of niche selection (54). Nevertheless, the effect of time on community composition indicated that large mesocosm communities were most influenced by demographic stochasticity arising from temporal influences. In this case, large mesocosms may more strongly reflect (i) random changes in births and deaths from a community that grew in number over time, (ii) stochasticity based on priority effects from slower time-delayed tracking, or (iii) may reflect sampling timepoints that underrepresented the larger total community.

Dispersal limitation as driver of metacommunity dynamics (considering all mesocosms at one time point) was present only in small sized mesocosms and suggests that multiple communities emerged from similar initial conditions in the small mesocosms. However, the interpretation of the dispersal limitation is ambiguous (e.g.(55)). It could be true dispersal limitation whereby niche spaces that are opened (i.e., when species become inactive in response to the initial salinity changes which increase habitat specialists (56) and/or the strong environmental changes) remain empty (57). However, it does not necessarily indicate true dispersal limitation between patches (55) or reduced immigration from a regional pool. Instead, it can be explained by the low richness of these mesocosm communities decreasing the likelihood that they contain superb dispersers. When dispersal rates are low, local adaptations to environmental fluctuations can lead to strengthened priority effects by preemptive taxa (58), which might have occurred during the experiment. For example, the many time-delayed associations between salinity and bacterial taxa in medium mesocosms could be a sign of transient priority effects where taxa (i.e., Bacteroidota and Proteobacteria but not Cyanobacteria) maintained abundances for a short period without environmental tracking. Nevertheless, with our data and the applied approaches, it is not possible to clearly support or exclude assembly processes and other factors that regulate them.

### Conclusions

Overall, our results partially align with those from previous studies which show that after disturbances, stochastic community assembly initially is important, but the dominant influence shifts to deterministic processes in later successional stages (e.g. (59)), especially when environmental conditions are stable. Dispersal limitation and ecological drift (demographic stochasticity) were drivers of metacommunity dynamics after community establishment with strong environmental fluctuations. Mesocosms with reduced environmental fluctuations may facilitate considerable time-delayed species sorting and thus possibly, a transient influence of priority effects. The novelty of our study is that we could show, by applying both path and network approaches, that the trajectories of (meta)community development are influenced by size-induced environmental fluctuations in concert with a salinity press disturbance. Our results represent the advantage of joining a network analysis together with metacommunity models, and stress that environmental fluctuations are important to consider in future community assembly studies as they can modify community assembly under natural conditions.

## Acknowledgements

This work was mainly supported by the Carl Tryggers Foundation. Sequencing was performed by the SNP&SEQ Technology Platform in Uppsala. The facility is part of the National Genomics Infrastructure (NGI) Sweden and Science for Life Laboratory. The SNP&SEQ Platform is also supported by the Swedish Research Council and the Knut and Alice Wallenberg Foundation. Computations were enabled by resources provided by the Swedish National Infrastructure for Computing (SNIC) at the Uppsala Multidisciplinary Center for Advanced Computational Science (UPPMAX) partially funded by the Swedish Research Council through grant agreement no. 2018-05973. This field mesocosm experiment has been made possible by the Swedish Infrastructure for Ecosystem Science (SITES), in this case the Erken Laboratory. SITES receives funding through the Swedish Research Council under the grant no. 2017-00635. This study was supported in part by resources and technical expertise from the Georgia Advanced Computing Resource Center, a partnership between the University of Georgia’s Office of the Vice President for Research and Office of the Vice President for Information Technology. The authors thank Christoffer Bergvall and Robin Hagblom for laboratory guidance and assistance, and Karsten Meier for help with fieldwork and data management.

## Data accessibility statement

The data supporting the results are archived in the public repository European Nucleotide Archive with accession number PRJEB26595 and environmental data are made available in the Swedish institutional repository, DiVA, (diva-portal.org) with the following accession number: diva2:1210995.

