## Supplementary Information for "Ecosystem size-induced environmental fluctuations affect the temporal dynamics of community assembly mechanisms"

**This file includes:**

Supplementary Methods and References

Supplementary Tables S2-S8

Supplementary Figures S1-S8

Supplementary Table S1 is supplied as a separate .xlxs file

### Supplementary Methods

*Experimental set-up and monitoring equipment*

The initial Lake Erken water measurements were as follows: 17.94 mg $L^{-1}$ total organic carbon (TOC), 0.671 mg $L^{-1}$ total nitrogen (TN), 0.016 mg $L^{-1}$ total phosphorus (TP), 25.86 relative fluorescence units (rfu) chlorophyll-*a* (chl-*a*), and 0.224 µm $L^{-1}$ colored fraction of dissolved organic matter (CDOM). The upper limit created for the mesocosm salinity, 6 ‰, corresponds to Baltic Sea salinity at Lake Erken latitude (1). Lake Erken is < 20 km from the coast and likely to have Baltic Sea bacteria in its water and sediment seed banks. This also entrains a likelihood that Baltic Sea bacteria dispersed through air and precipitation into the mesocosms.

Water sample monitoring was conducted for conductivity and temperature (Cond 3210 conductivity meter Xylem Analytics Germany Sales GmbH & Co. KG, Weilheim, Germany), and depth integrated pH (micropH 2001 Crison Instruments, S.A., Alella, Spain), chlorophyll-*a*, and CDOM fluorescence (Aqua Fluor Handheld Fluorometer/Turbidimeter Turner Designs, San Jose, CA, USA at 395 nm and 350 nm, respectively).

*Bacterial community composition of external sources*

Ongoing immigration from external sources was characterized using sterile air and rain traps (n = 3, SA = 96 $cm^2$) interspersed beside the mesocosm units. Sheltered air traps contained 200 mL sterile filtered water; rain traps were uncovered. Every 8 days, trap samples were collected, pooled, and processed with mesocosm samples for 16S rRNA gene amplicon sequencing.

*Molecular sample preparation*

Samples for RNA analysis were treated with DNase I (Invitrogen), checked using 35-cycle PCR amplification, and then transcribed to cDNA (2). Primers 341F (3) and 805RN (4) containing Illumina adaptors (5) were used for amplification. A second-step PCR barcoded samples. Purified samples were quantified with Quant-iT PicoGreen dsDNA Reagent Kit (Invitrogen, Carlsbad, CA, USA), combined in equimolar amounts in two pools. Each pool was gel purified (GeneJET gel extraction, ThermoFisher Scientific, Uppsala, Sweden). See DOI for a detailed protocol: dx.doi.org/10.17504/protocols.io.xekfjcw

*Additional data processing details*

Sequences were trimmed to 280 bp and 200 bp for forward and reverse reads, respectively, and merged with at most two expected errors. Following dereplication and chimeric sequence removal from the Amplicon Sequence Variants (ASVs), the SILVA v. 138.1 reference database (6) (August 2020) was used to assign taxonomy at 99 % threshold. ASVs not identified as Bacteria were removed. Forty ASVs identified as extraction or PCR contaminants using R package "decontam" v 1.8.0 ref. (7) with default settings using the prevalence-based approach were removed from 11 samples leaving 12 068 ASVs.

*Network analysis details*

For equal time windows between samplings (8-days), samples from day 2 and 4 were removed. Samples from day 64 were also removed due to the low sequence retention (Table S2). The 50 most abundant ASVs from each size category that were used comprised, on average, 70 % (range 53–76 %), 72 % (range 59–80 %), and 63 % (range 47–72 %) of the relative bacterial abundance in small, medium, and large mesocosms, respectively. 'Simple' method (simple average method) was used to summarize replicate data, and p-values for pairwise LS correlations were determined using the ‘mixed’ approach (8). LS values were considered statistically significant if $p \leq 0.01$ and Q (the false discovery rate) $\leq 0.01$.

**Table S1.** Statistics of sequences from DADA2 sequence processing. (See Excel file)

**Table S2.** Samples not meeting the 5028 sequence per sample evenness requirement for inclusion in beta diversity analysis.

| Sample Day | Mesocosm ID | Total Sequences |
|---|---|---|
| D16 | 19 | 0 |
| D56 | 13 | 5 |
| D48 | 29 | 752 |
| D24 | 2 | 1993 |
| D64 | 33 | 2067 |
| D64 | 48 | 2276 |
| D64 | 45 | 2527 |
| D64 | 31 | 2546 |
| D64 | 6 | 2793 |
| D64 | 12 | 2952 |
| D64 | 37 | 3064 |
| D64 | 39 | 3189 |
| D64 | 17 | 3353 |
| D64 | 44 | 3541 |
| D64 | Rain | 3783 |
| D64 | 36 | 3900 |
| D64 | 5 | 3939 |
| D64 | 40 | 4106 |
| D56 | Air | 4137 |
| D64 | 15 | 4161 |
| D64 | Air | 4401 |
| D64 | 27 | 4479 |
| D64 | 47 | 4626 |
| D32 | 1 | 4666 |
| D64 | 43 | 4670 |
| D64 | 16 | 4770 |
| D64 | 35 | 4982 |
| D64 | 11 | 4996 |

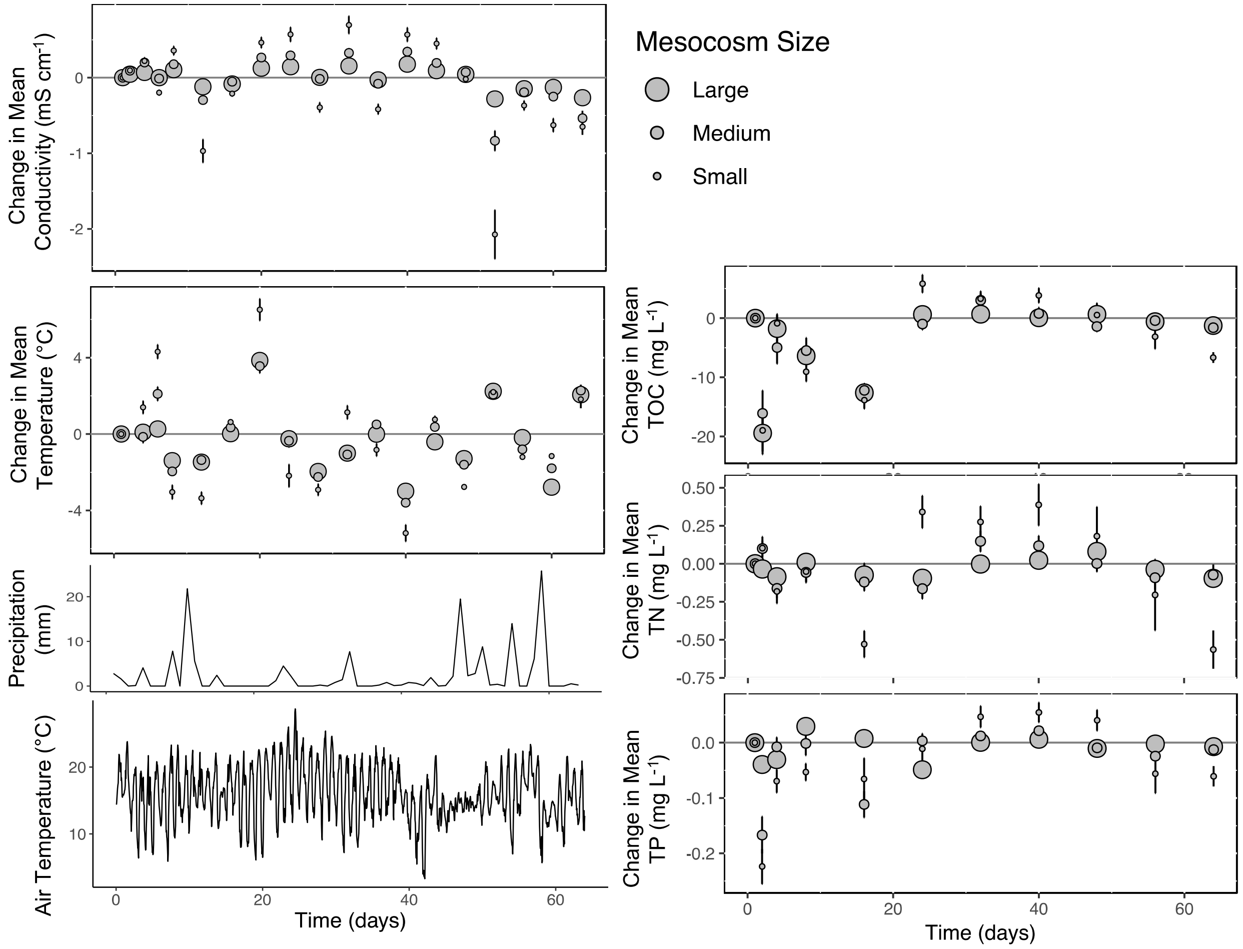


**Figure S1.** Change in mean conductivity, water temperature, and nutrients of experimental mesocosms from previous sampling day. Daily precipitation and air temperature were gathered from Svanberga station, Sweden, SMHI. Error bars represent standard error from the mean. (n = 16)

**Table S3.** Results from nonparametric tests with an ANOVA-type statistic for repeated measures to compare changes in environmental variables between mesocosm sizes.

| Environmental Variable (Absolute Change) | Category | Term | Statistic | df | p-value |
|---|---|---|---|---|---|
| Conductivity (mS/cm) | Overall | Size | 7.93 | 1.9 | **0.001** |
| | | Day | 86.52 | 3.4 | **>0.001** |
| | | Size:Day | 5.15 | 5.5 | **>0.001** |
| | Small vs Medium | Size | 2.99 | 1.0 | 0.084 |
| | Small vs Large | Size | 15.97 | 1.0 | **>0.001** |
| | Medium vs Large | Size | 5.50 | 1.0 | **0.019** |
| Temperature (°C) | Overall | Size | 108.35 | 1.7 | **>0.001** |
| | | Day | 91.84 | 4.7 | **>0.001** |
| | | Size:Day | 11.75 | 6.4 | **>0.001** |
| | Small vs Medium | Size | 74.55 | 1.0 | **>0.001** |
| | Small vs Large | Size | 190.41 | 1.0 | **>0.001** |
| | Medium vs Large | Size | 33.15 | 1.0 | **>0.001** |
| Chlorophyll-a (RFU) | Overall | Size | 7.23 | 1.5 | **0.002** |
| | | Day | 19.94 | 10.3 | **>0.001** |
| | | Size:Day | 2.66 | 16.2 | **>0.001** |
| | Small vs Medium | Size | 3.83 | 1.0 | 0.050 |
| | Small vs Large | Size | 12.28 | 1.0 | **<0.001** |
| | Medium vs Large | Size | 4.66 | 1.0 | **0.031** |
| CDOM (mg/L) | Overall | Size | 32.68 | 1.6 | **<0.001** |
| | | Day | 26.69 | 10.7 | **<0.001** |
| | | Size:Day | 6.27 | 16.7 | **<0.001** |
| | Small vs Medium | Size | 22.84 | 1.0 | **<0.001** |
| | Small vs Large | Size | 48.72 | 1.0 | **<0.001** |
| | Medium vs Large | Size | 14.59 | 1.0 | **<0.001** |
| pH | Overall | Size | 4.13 | 1.9 | **0.019** |
| | | Day | 0.59 | 4.4 | 0.688 |
| | | Size:Day | 3.34 | 8.0 | **0.001** |
| | Small vs Medium | Size | 0.95 | 1.0 | 0.329 |
| | Small vs Large | Size | 3.72 | 1.0 | 0.054 |
| | Medium vs Large | Size | 6.39 | 1.0 | **0.012** |
| Cell Abundance (cells/mL) | Overall | Size | 16.81 | 2.0 | **<0.001** |
| | | Day | 26.37 | 10.1 | **<0.001** |
| | | Size:Day | 3.69 | 16.5 | **<0.001** |
| | Small vs Medium | Size | 1.83 | 1.0 | 0.177 |
| | Small vs Large | Size | 31.61 | 1.0 | **<0.001** |
| | Medium vs Large | Size | 18.75 | 1.0 | **<0.001** |
| TOC (mg/L) | Overall | Size | 16.76 | 1.9 | **<0.001** |
| | | Day | 30.35 | 6.5 | **<0.001** |
| | | Size:Day | 2.02 | 11.1 | **0.022** |
| | Small vs Medium | Size | 4.87 | 1.0 | **0.027** |
| | Small vs Large | Size | 32.34 | 1.0 | **<0.001** |
| | Medium vs Large | Size | 14.68 | 1.0 | **<0.001** |
| TN (mg/L) | Overall | Size | 41.10 | 1.9 | **<0.001** |
| | | Day | 3.11 | 7.4 | **0.002** |
| | | Size:Day | 2.07 | 12.3 | **0.015** |
| | Small vs Medium | Size | 18.84 | 1.0 | **<0.001** |
| | Small vs Large | Size | 87.81 | 1.0 | **<0.001** |
| | Medium vs Large | Size | 21.50 | 1.0 | **<0.001** |
| TP (mg/L) | Overall | Size | 24.25 | 1.8 | **<0.001** |
| | | Day | 20.27 | 7.2 | **<0.001** |
| | | Size:Day | 1.24 | 12.1 | 0.245 |
| | Small vs Medium | Size | 10.53 | 1.0 | **0.001** |
| | Small vs Large | Size | 66.49 | 1.0 | **<0.001** |
| | Medium vs Large | Size | 10.93 | 1.0 | **0.001** |

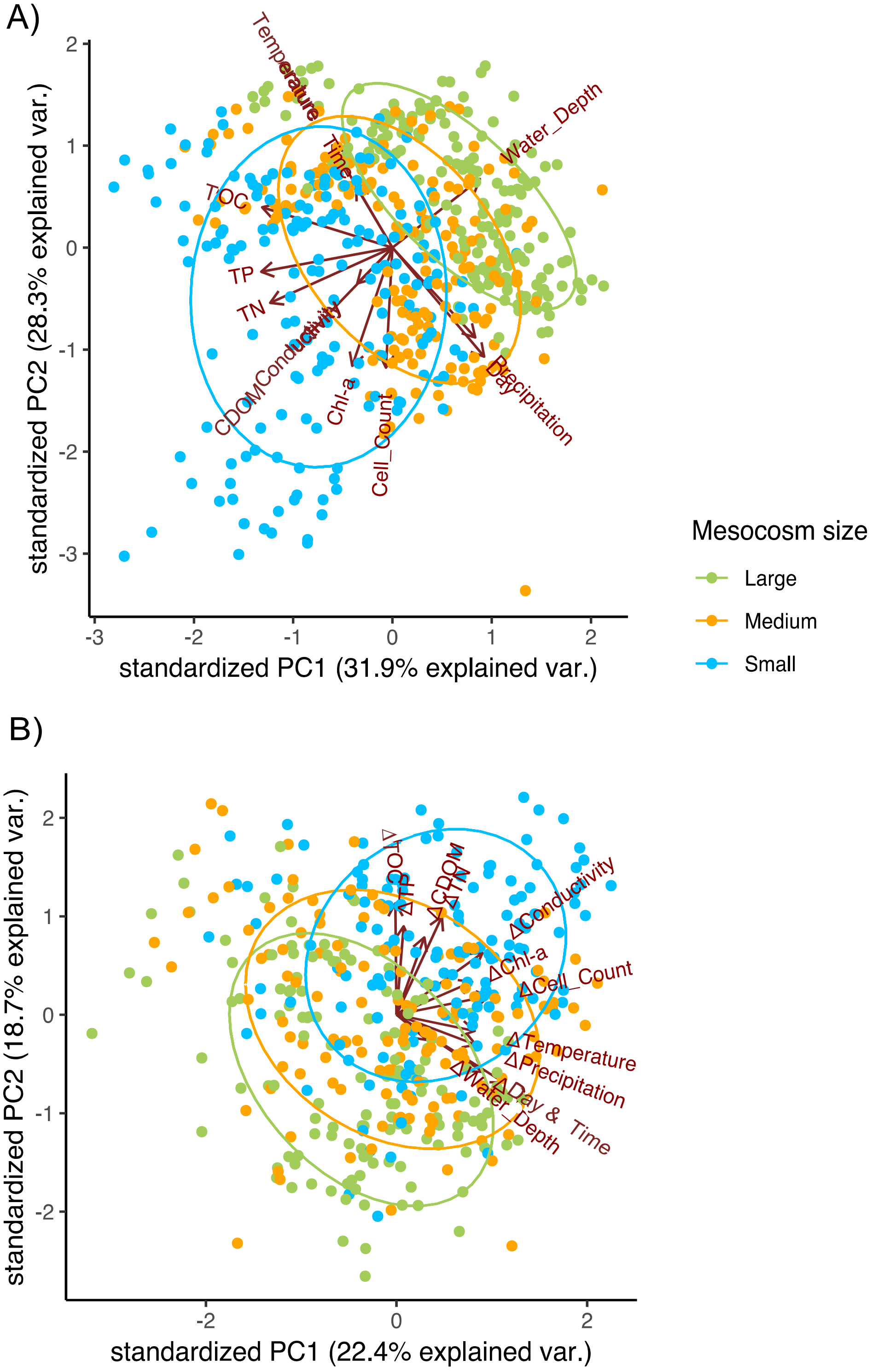


**Figure S2.** Principal components analysis plots of log transformed environmental variables using raw data (A) or absolute changes from sampling day-to-sampling day (B). Different colors show mesocosm size. Environmental vector correlations shown were significant at $p < 0.05$.

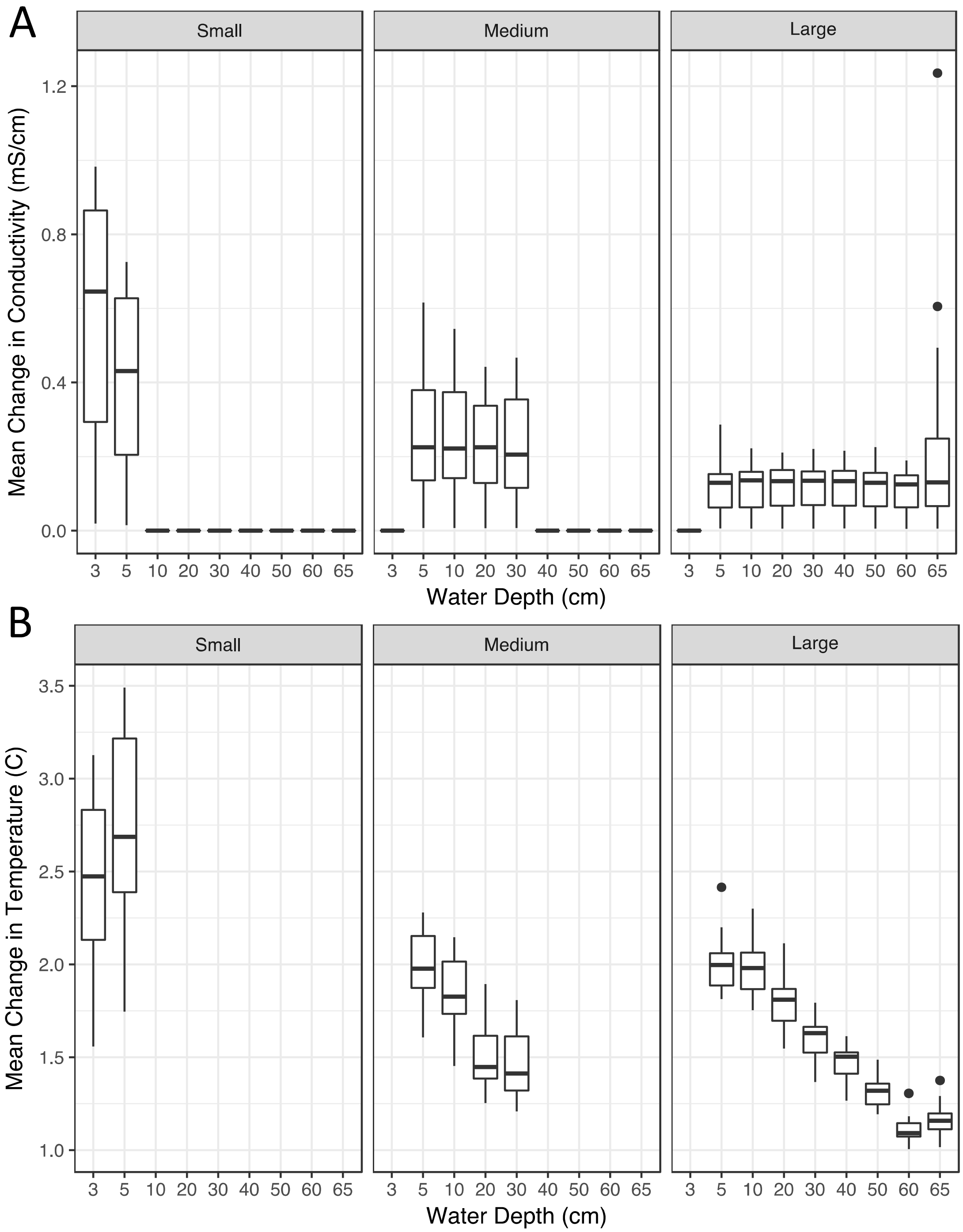


**Figure S3.** Depth profiles for mean of changes in environmental variables (A: conductivity, B: temperature) over experiment duration (64 days) in mesocosms of different size. (n = 16)

A)

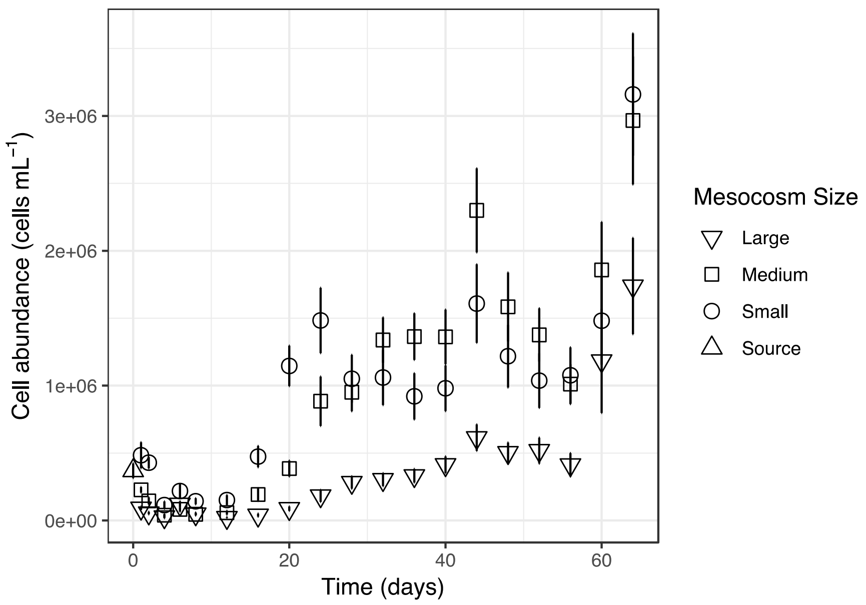


B)

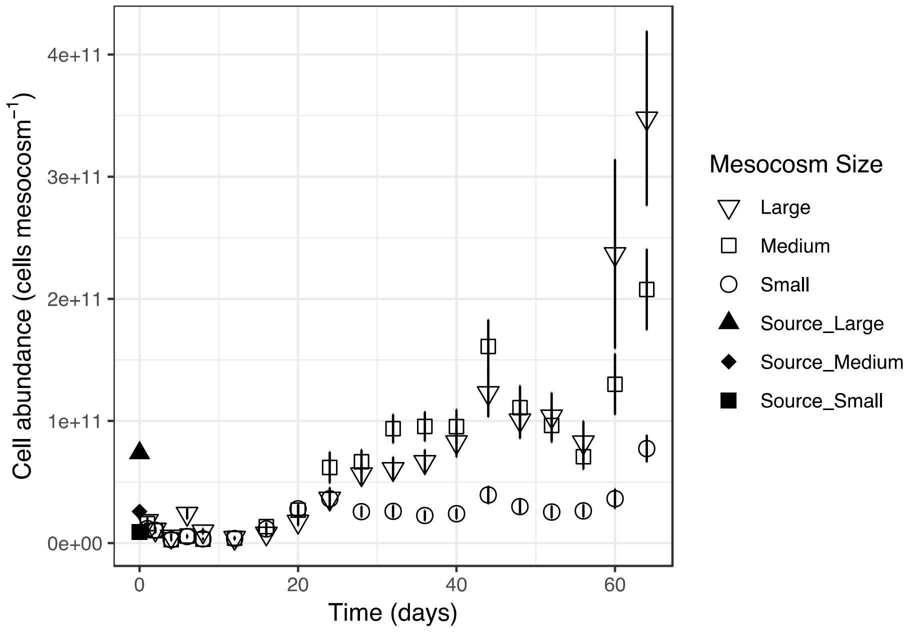


**Figure S4.** Cell abundance as A: mean cells $mL^{-1}$ and B: total cells per mesocosm. Cell abundance from lake source at Day 0. Error bars are standard error. (n = 16)

A)

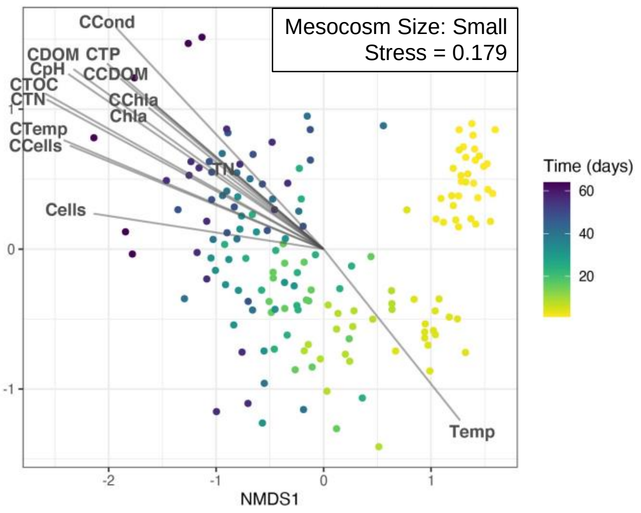

Mesocosm Size: Small
Stress = 0.179
CCond
CDOM CTP
CpH
CCDOM
CTOC
CTN
CChla
Chla
CTemp
CCells
TN
Cells
Temp
Time (days)
60
40
20
NMDS2
NMDS1
-2
-1
0
1


B)

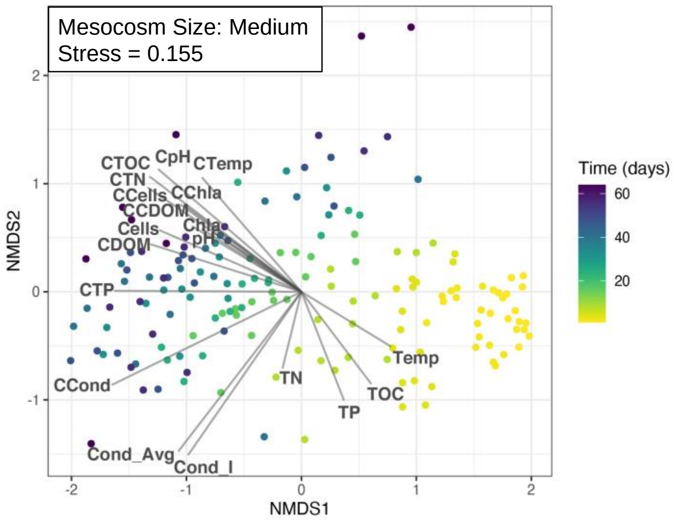

Mesocosm Size: Medium
Stress = 0.155
CTOC
CpH
CTemp
CTN
CCells
CChla
CCDOM
Chla
Cells
pH
CDOM
CTP
CCond
TN
TOC
TP
Temp
Cond_Avg
Cond_I
Time (days)
60
40
20
NMDS2
NMDS1
-2
-1
0
1
2


C)

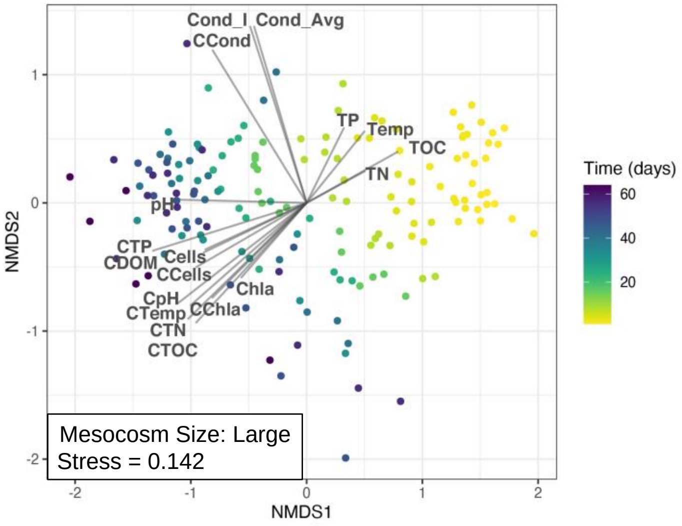

Cond_I
Cond_Avg
CCond
TP
Temp
TOC
TN
pH
CTP
CDOM
Cells
CCells
Chla
CpH
CTemp
CChla
CTN
CTOC
Mesocosm Size: Large
Stress = 0.142
Time (days)
60
40
20
NMDS2
NMDS1
-2
-1
0
1
2

D)

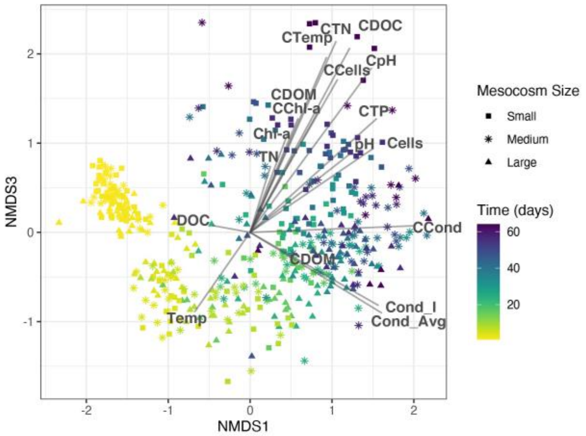
CTemp
CTN
CDOC
CCells
CpH
CDOM
CChl-a
CTP
Chl-a
pH
Cells
TN
DOC
CCond
CDOM
Cond_I
Cond_Avg
Temp
NMDS1
NMDS3
Mesocosm Size
Small
Medium
Large
Time (days)
60
40
20

G)

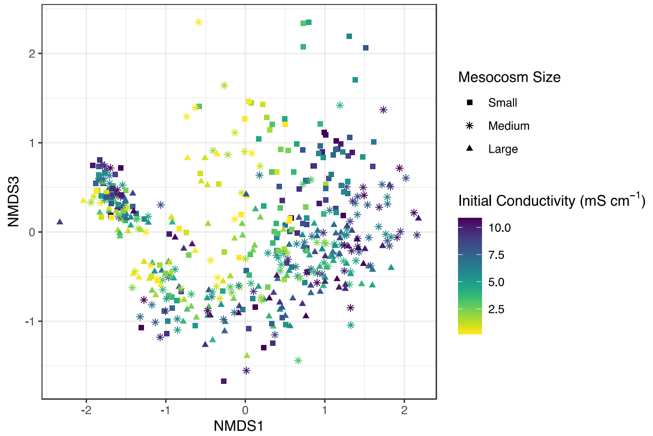
NMDS1
NMDS3
Mesocosm Size
Small
Medium
Large
Initial Conductivity (mS cm$^{-1}$)
10.0
7.5
5.0
2.5

E)

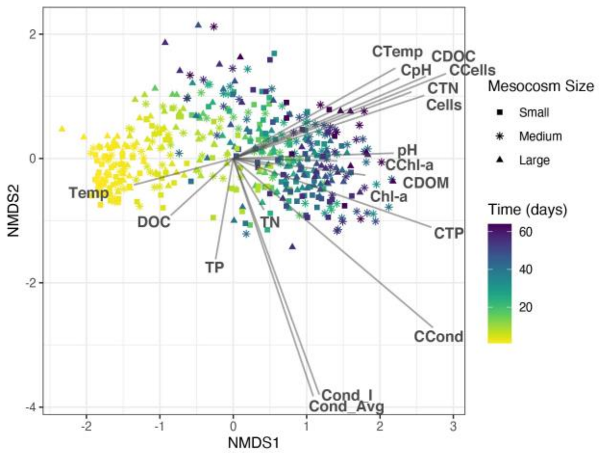
CTemp
CDOC
CpH
CCells
CTN
Cells
pH
CChl-a
CDOM
Chl-a
CTP
Temp
DOC
TN
TP
CCond
Cond_I
Cond_Avg
NMDS1
NMDS2
Mesocosm Size
Small
Medium
Large
Time (days)
60
40
20

H)

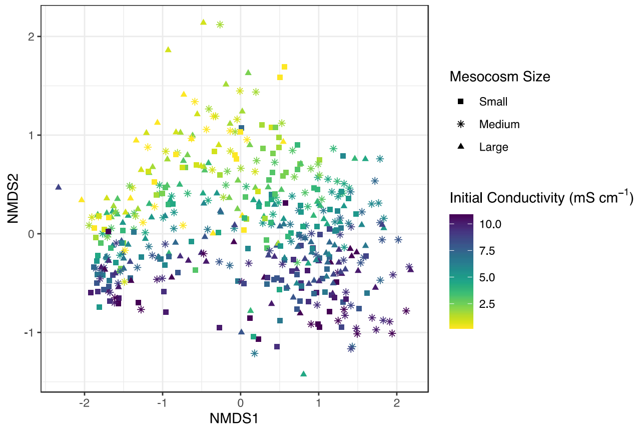
NMDS1
NMDS2
Mesocosm Size
Small
Medium
Large
Initial Conductivity (mS cm$^{-1}$)
10.0
7.5
5.0
2.5

F)

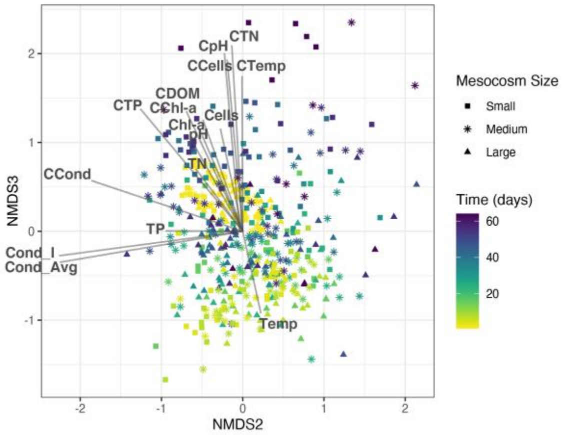
CpH
CTN
CCells
CTemp
CDOM
CTP
CChl-a
Cells
Chl-a
pH
TN
CCond
TP
Cond_I
Cond_Avg
Temp
NMDS2
NMDS3
Mesocosm Size
Small
Medium
Large
Time (days)
60
40
20

I)

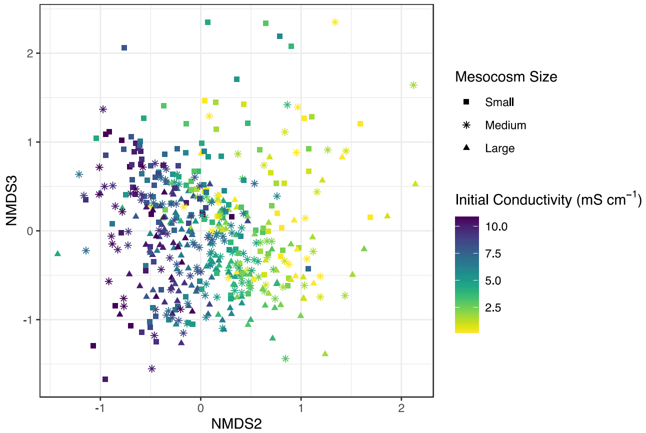
NMDS2
NMDS3
Mesocosm Size
Small
Medium
Large
Initial Conductivity (mS cm$^{-1}$)
10.0
7.5
5.0
2.5

**Figure S5.** Nonmetric multidimensional scaling graphs of Bray-Curtis dissimilarity matrices for bacterial communities in A) small, B) medium, and C) large mesocosm sizes and in all mesocosm sizes with environmental vectors and points colored by time (D-F) or by initial conductivity (G-I). 'C' in environmental vector names indicates cumulative change. When applicable, environmental vectors were depth averaged values. Cond_I signifies initial mesocosm conductivity from conductivity gradient. NMDS plots repeated to show all combinations of two of the three axes. All environmental vectors $p < 0.05$. NMDS stress = 0.128.

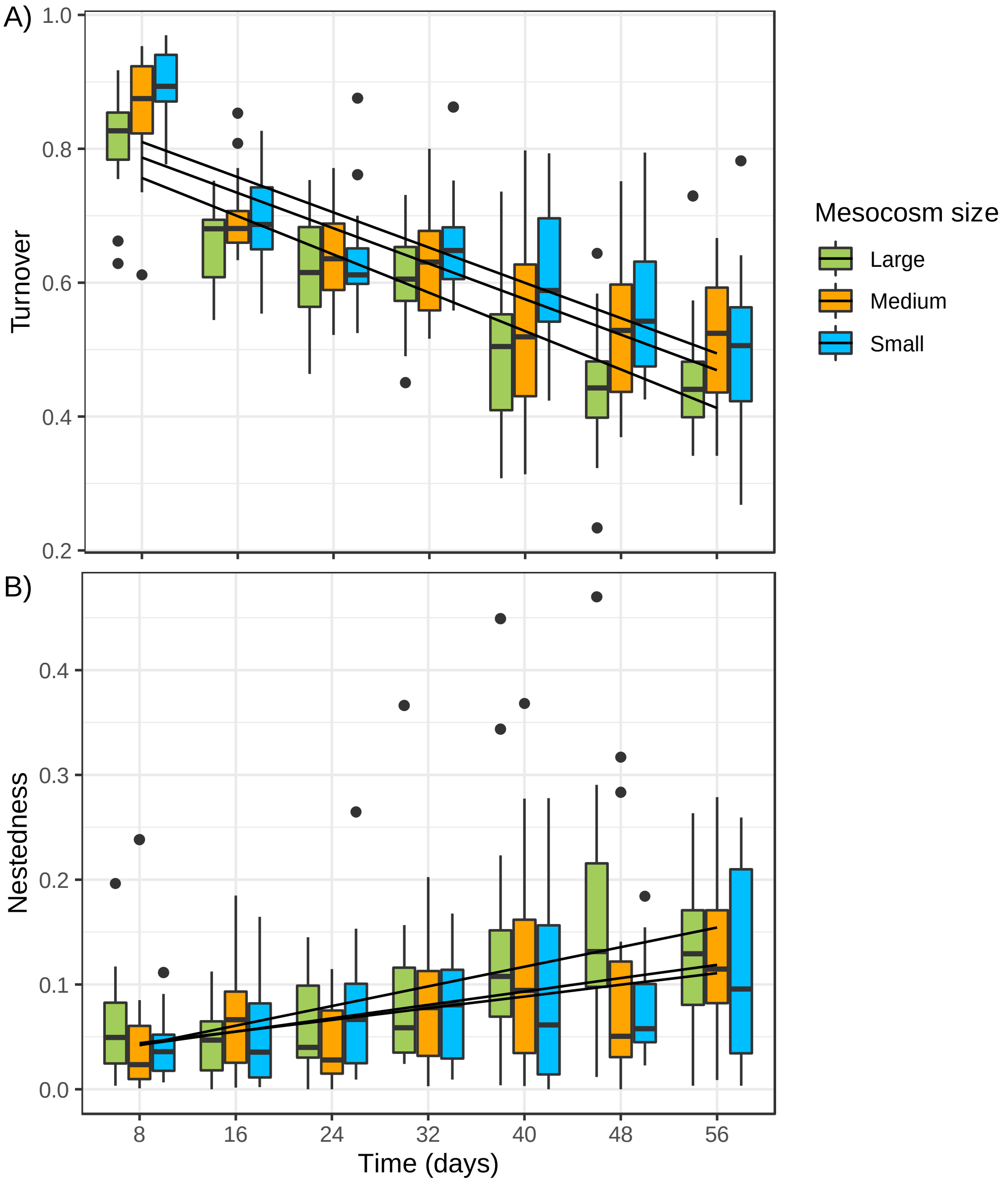


**Figure S6.** Beta diversity in different mesocosm sizes over time compared to the prior time point based on Jaccard dissimilarity index of taxa presence-absence and partitioned into taxa turnover (A) and nestedness (B). (n = 16)

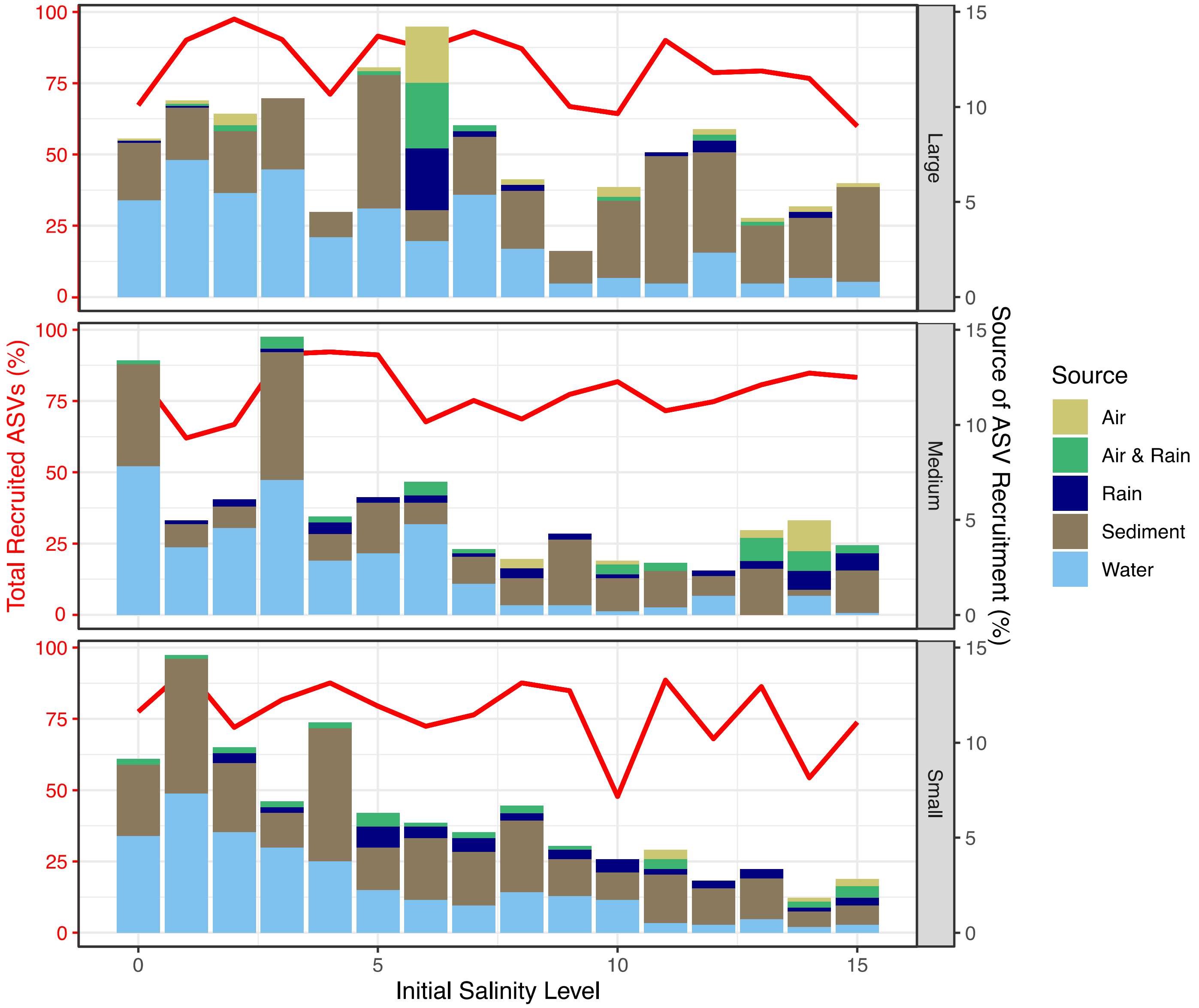


**Figure S7.** The percent of unique ASVs recruited over the duration of the experiment in each mesocosm at different levels of initial salinity disturbance with level 0 representing 0 ‰ salinity and level 15 representing 6 ‰ salinity. Panels are ordered by mesocosm size with large mesocosms in the top panel. The line plot represents total unique ASV recruitment including from unknown sources. The bar plot represents ASVs recruited from identified sources with the total height of the bar showing the percentage of ASVs recruited to that mesocosm from an identified source. Air and rain had 156 ASVs recruited in common.

**Table S4**: Standardized estimates for the path analysis of the small mesocosms. Significant effects are with a Benjamini-Hochberg correction. Standardized Root Mean Square Residual = 0.229.

| **Small Mesocosms** | | | | | | | | |
|---|---|---|---|---|---|---|---|---|
| Environmental and Community Variables | | Cond | Temp | Chl-*a* | CDOM | TOC | TN | TP |
| $\beta_{bc} \leftarrow \langle J \rangle$ | -0.173*** | | | | | | | |
| $\beta_{bc} \leftarrow \Delta S$ | 0.028 | | | | | | | |
| $\beta_{bc} \leftarrow \Delta t$ | 0.326*** | | | | | | | |
| $\beta_{bc} \leftarrow \Delta x$ | 0.076*** | | | | | | | |
| $\beta_{bc} \leftarrow \Delta E$ | 0.925[‡] | 0.320*** | 0.046* | 0.226*** | -0.056 | -0.122*** | 0.140*** | 0.015 |
| $\Delta S \leftarrow \Delta J$ | 0.016 | | | | | | | |
| $\Delta E \leftarrow \Delta t$ | 1.201[‡] | -0.022 | 0.558*** | 0.160*** | 0.312*** | 0.014 | 0.096** | 0.039 |
| $\Delta E \leftarrow \Delta x$ | | 0.052* | | | | | | |

Cond = conductivity, Temp = temperature, Chl-*a* = chlorophyll-*a*
* p < 0.05 to 0.01, ** p 0.01 to 0.001, *** p <0.001
[‡] Total of the absolute values of standardized estimates in the row

**Table S5**: Standardized estimates for the path analysis of the medium mesocosms. Significant effects are with a Benjamini-Hochberg correction. Standardized Root Mean Square Residual = 0.127.

| **Medium Mesocosms** | | | | | | | | |
|---|---|---|---|---|---|---|---|---|
| Environmental and Community Variables | | Cond | Temp | Chl-*a* | CDOM | TOC | TN | TP |
| $\beta_{bc} \leftarrow \langle J \rangle$ | -0.133* | | | | | | | |
| $\beta_{bc} \leftarrow \Delta S$ | 0.084* | | | | | | | |
| $\beta_{bc} \leftarrow \Delta t$ | 0.273*** | | | | | | | |
| $\beta_{bc} \leftarrow \Delta x$ | 0.013 | | | | | | | |
| $\beta_{bc} \leftarrow \Delta E$ | 0.773[‡] | 0.511*** | -0.016 | -0.072 | 0.083 | -0.009 | 0.057 | 0.025 |
| $\Delta S \leftarrow \Delta J$ | 0.227*** | | | | | | | |
| $\Delta E \leftarrow \Delta t$ | 1.477[‡] | -0.026 | 0.551*** | 0.218*** | 0.284*** | 0.235*** | -0.014 | 0.149*** |
| $\Delta E \leftarrow \Delta x$ | | 0.283*** | | | | | | |

Cond = conductivity, Temp = temperature, Chl-*a* = chlorophyll-*a*
* p < 0.05 to 0.01, ** p 0.01 to 0.001, *** p <0.001
[‡] Total of the absolute values of standardized estimates in the row

**Table S6**: Standardized estimates for the path analysis of the large mesocosms. Significant effects are with a Benjamini-Hochberg correction. Standardized Root Mean Square Residual = 0.116.

| **Large Mesocosms** | | | | | | | | |
|---|---|---|---|---|---|---|---|---|
| Environmental and Community Variables | | Cond | Temp | Chl-*a* | CDOM | TOC | TN | TP |
| $\beta_{bc} \leftarrow \langle J \rangle$ | -0.136* | | | | | | | |
| $\beta_{bc} \leftarrow \Delta S$ | 0.174*** | | | | | | | |
| $\beta_{bc} \leftarrow \Delta t$ | 0.370*** | | | | | | | |
| $\beta_{bc} \leftarrow \Delta x$ | -0.058* | | | | | | | |
| $\beta_{bc} \leftarrow \Delta E$ | 0.766[‡] | 0.427*** | -0.084*** | 0.099* | 0.075 | -0.038 | -0.007 | -0.036 |
| $\Delta S \leftarrow \Delta J$ | 0.147*** | | | | | | | |
| $\Delta E \leftarrow \Delta t$ | 1.579[‡] | -0.014 | 0.485*** | 0.124*** | 0.349*** | 0.308*** | 0.099*** | 0.200** |
| $\Delta E \leftarrow \Delta x$ | | 0.298*** | | | | | | |

Cond = conductivity, Temp = temperature, Chl-*a* = chlorophyll-*a*
* p < 0.05 to 0.01, ** p 0.01 to 0.001, *** p <0.001
[‡] Total of the absolute values of standardized estimates in the row

**Table S7**: Summary statistics for the constructed complete (Bacteria + Env) and bacterial subnetworks of 50 most abundant Bacteria (ASVs) and environmental parameters of small, medium, and large mesocosms (Figure 4). Note that only the results of the significant associations are listed.

| | Small | | Medium | | Large | |
|---|---|---|---|---|---|---|
| **Parameters** | **Bacteria** | **Bacteria + Env** | **Bacteria** | **Bacteria + Env** | **Bacteria** | **Bacteria + Env** |
| Nodes (without edge to environmental variable) | 37 | 43 (18) | 49 | 58 (4) | 50 | 60 (6) |
| Edges | 44 | 73 | 246 | 385 | 293 | 454 |
| Positive-delayed edges (% of all edges) | 5 (11.4%) | 5 (6.8%) | 33 (13.4%) | 57 (14.8%) | 32 (10.9%) | 54 (11.9%) |
| Negative-delayed edges (% of all edges) | 4 (9.1%) | 10 (13.7%) | 34 (13.8%) | 71 (18.4%) | 37 (12.6%) | 89 (19.6%) |
| Diameter (radius) | 7 (4) | 6 (3) | 6 (3) | 5 (3) | 8 (4) | 6 (3) |
| Delayed/non-delayed associations | 9 / 35 | 15 / 58 | 67 / 179 | 128 / 257 | 69 / 224 | 143 / 311 |
| *Connectivity* | | | | | | |
| Average number of neighbors | 3.08 | 4.21 | 10.04 | 13.28 | 11.72 | 15.13 |
| Network density | 0.134 | 0.156 | 0.209 | 0.23 | 0.239 | 0.256 |
| *Likelihood for uneven distribution of edges* | | | | | | |
| Network heterogeneity | 0.537 | 0.635 | 0.619 | 0.54 | 0.527 | 0.55 |
| Network centralization | 0.138 | 0.271 | 0.281 | 0.249 | 0.219 | 0.524 |
| *Identifying small-world properties* | | | | | | |
| Characteristic path length | 3.014 | 2.643 | 2.395 | 2.13 | 2.781 | 2.128 |
| Clustering coefficient | 0.353 | 0.451 | 0.589 | 0.60 | 0.513 | 0.537 |

Env = environmental parameter

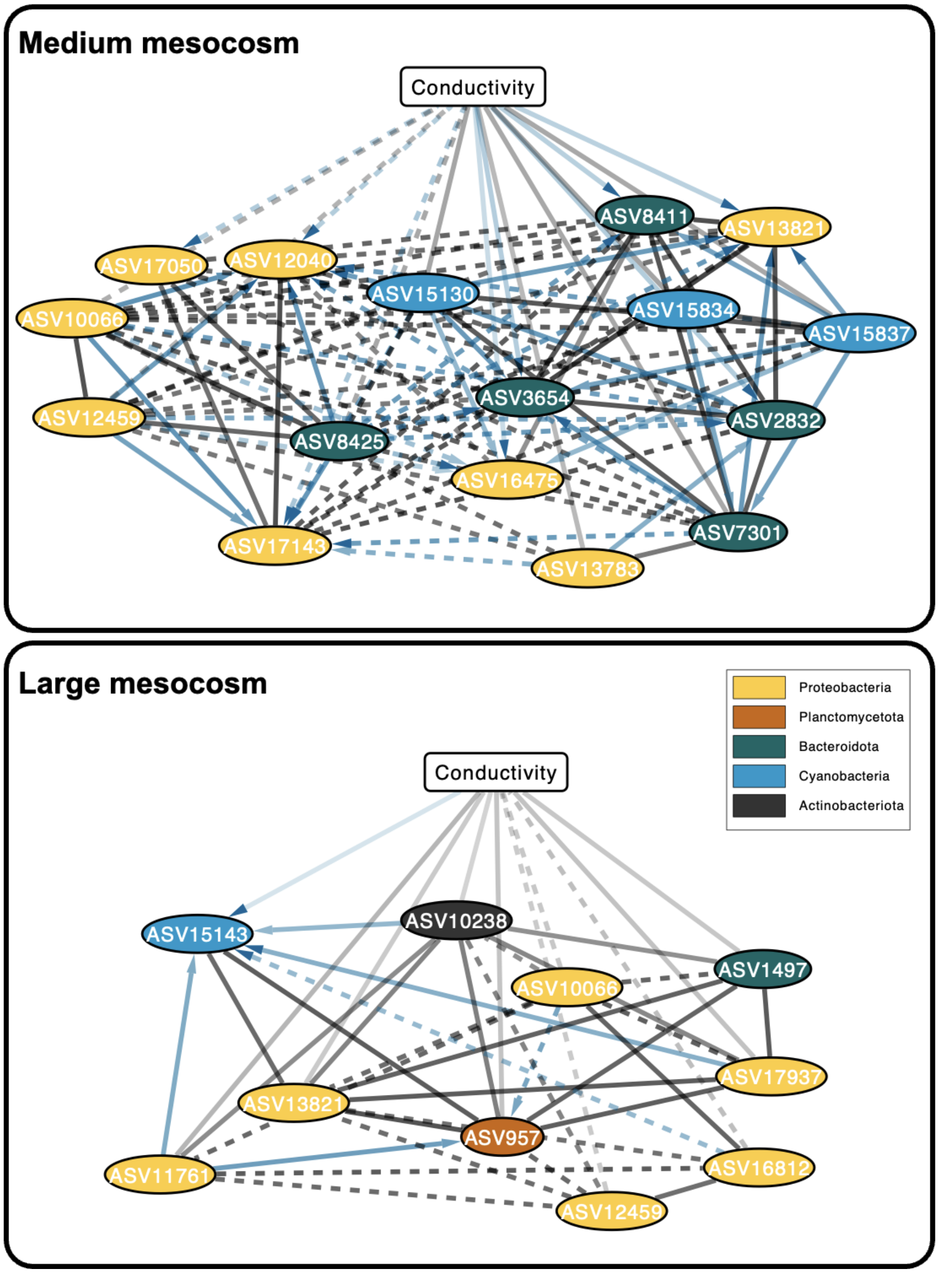


**Figure S8.** Subsets of the association networks of the mesocosm size categories (n = 16), highlighting bacterial ASVs that had association with conductivity. All significant ($p \leq 0.01$ and $Q \leq 0.01$) pairwise local similarity (LS) correlations $\geq 0.05$ are shown as edges in the networks. Each node represents an ASV (ellipse) or an environmental factor (rectangle). Edge transparency is proportional to the association strength (based on LS values). Solid lines refer to positive associations while dashed lines to negative ones. Edge colors indicate delayed (blue) and non-delayed (black) associations between ASVs and/or environmental variables. Arrows point toward the lagging node. Note that a network graph of small mesocosms is not present due to the lack of any significant correlation.

**Table S8**: Summary of the aquatic bacterial subnetworks in small, medium, and large mesocosms with relation to water conductivity over time.

| Parameters | Small | Medium | Large |
|---|---|---|---|
| Nodes | 1 | 17 | 11 |
| Edges | - | 100 | 42 |
| Positive edges between ASV and Conductivity | - | 10 | 7 |
| Negative edges between ASV and Conductivity | - | 6 | 3 |
| Delayed/non-delayed associations between ASVs and Conductivity | - | 8 / 8 | 1 / 9 |
