## Supplementary Table S1 for "Ecosystem size-induced environmental fluctuations affect the temporal dynamics of community assembly mechanisms"

Supplementary Table S1. Statistics of sequences from DADA2 sequence processing.

| sample name | input | filtered | merged | nonchimeric | contaminants removed | bacteria only | sample coverage |
|---|---|---|---|---|---|---|---|
| D0MSDNAMa | 145044 | 108836 | 59497 | 55639 | 51135 | 51138 | 0.9997 |
| D0MSDNAMb | 140913 | 105305 | 55148 | 51684 | 49487 | 49487 | 0.9997 |
| D0MSDNAMc | 129822 | 95639 | 50100 | 46609 | 42052 | 42052 | 0.9997 |
| D0MWDNAMa | 129072 | 100555 | 70873 | 52742 | 45800 | 45800 | 0.9999 |
| D0MWDNAMb | 126110 | 97781 | 73497 | 57193 | 44883 | 44883 | 0.9999 |
| D0MWDNAMc | 133015 | 103571 | 77518 | 59474 | 55704 | 55704 | 1.0000 |
| D1M1 | 33712 | 27548 | 19752 | 15740 | 15683 | 15683 | 0.9998 |
| D1M10 | 34654 | 27656 | 20402 | 16578 | 16570 | 16570 | 0.9998 |
| D1M11 | 28609 | 22867 | 15783 | 13718 | 13655 | 13655 | 1.0000 |
| D1M12 | 42982 | 34460 | 25766 | 19999 | 19994 | 19994 | 0.9998 |
| D1M13 | 30843 | 24929 | 19466 | 15841 | 15841 | 15841 | 0.9999 |
| D1M14 | 50858 | 40847 | 29798 | 21955 | 21904 | 21918 | 0.9997 |
| D1M15 | 25793 | 20508 | 12890 | 11216 | 11025 | 11025 | 0.9999 |
| D1M16 | 44810 | 35861 | 28846 | 26367 | 26299 | 26299 | 0.9999 |
| D1M17 | 34398 | 28241 | 22429 | 19330 | 19330 | 19330 | 0.9999 |
| D1M18 | 76837 | 62127 | 48619 | 37384 | 37358 | 37358 | 0.9998 |
| D1M19 | 55562 | 45521 | 37106 | 30440 | 30422 | 30422 | 0.9999 |
| D1M2 | 39111 | 31600 | 23602 | 20020 | 19739 | 19739 | 0.9998 |
| D1M20 | 54390 | 44131 | 31469 | 26156 | 26000 | 26000 | 1.0000 |
| D1M21 | 41792 | 34172 | 24206 | 19670 | 19633 | 19633 | 0.9996 |
| D1M22 | 30015 | 24555 | 18212 | 14527 | 14522 | 14522 | 0.9999 |
| D1M23 | 40065 | 32102 | 19475 | 16308 | 16057 | 16057 | 0.9999 |
| D1M24 | 32796 | 26400 | 19577 | 17394 | 17311 | 17311 | 0.9999 |
| D1M25 | 36328 | 28651 | 17461 | 13117 | 13093 | 13093 | 0.9997 |
| D1M26 | 64951 | 51697 | 39150 | 30162 | 30142 | 30142 | 0.9999 |
| D1M27 | 38019 | 30352 | 23848 | 22249 | 22081 | 22203 | 0.9998 |
| D1M28 | 36654 | 29317 | 21277 | 19101 | 18966 | 18966 | 0.9999 |
| D1M29 | 74871 | 60057 | 43109 | 33308 | 33264 | 33264 | 0.9997 |
| D1M3 | 52023 | 42312 | 33050 | 27182 | 27051 | 27051 | 0.9999 |
| D1M30 | 46813 | 37178 | 25503 | 22740 | 22425 | 22425 | 0.9998 |
| D1M31 | 59000 | 46774 | 39040 | 38772 | 36584 | 36610 | 0.9999 |
| D1M32 | 56395 | 44452 | 37289 | 36756 | 36324 | 36345 | 0.9999 |
| D1M33 | 55692 | 44626 | 27826 | 23988 | 23056 | 23056 | 0.9999 |
| D1M34 | 61486 | 49685 | 37591 | 29987 | 29965 | 29965 | 0.9999 |
| D1M35 | 80491 | 65744 | 56342 | 53393 | 53240 | 53277 | 0.9999 |
| D1M36 | 49124 | 39929 | 27911 | 23371 | 23275 | 23275 | 0.9998 |
| D1M37 | 62896 | 51790 | 45942 | 44836 | 44776 | 44836 | 1.0000 |
| D1M38 | 85160 | 69431 | 54674 | 44436 | 44336 | 44336 | 0.9999 |
| D1M39 | 61970 | 50454 | 39123 | 32198 | 32187 | 32187 | 0.9998 |
| D1M4 | 55355 | 45293 | 35037 | 27668 | 27619 | 27619 | 0.9999 |

| | | | | | | | |
|---|---|---|---|---|---|---|---|
| D1M40 | 69549 | 55793 | 47943 | 47649 | 46628 | 46699 | 0.9999 |
| D1M41 | 70471 | 57579 | 43412 | 35071 | 34983 | 34983 | 0.9999 |
| D1M42 | 64344 | 50703 | 37834 | 34402 | 33918 | 34177 | 0.9998 |
| D1M43 | 65198 | 53040 | 45151 | 43065 | 42922 | 43043 | 0.9999 |
| D1M44 | 98238 | 79806 | 68724 | 64538 | 64383 | 64473 | 0.9999 |
| D1M45 | 45423 | 36733 | 24530 | 20716 | 20479 | 20479 | 0.9998 |
| D1M46 | 79754 | 64933 | 54748 | 49011 | 48500 | 48924 | 1.0000 |
| D1M47 | 52720 | 43095 | 37829 | 36110 | 36023 | 36030 | 0.9999 |
| D1M48 | 28715 | 22818 | 14142 | 11913 | 11788 | 11788 | 0.9998 |
| D1M5 | 30045 | 24624 | 19099 | 15584 | 15584 | 15584 | 0.9998 |
| D1M6 | 65610 | 53068 | 40112 | 30599 | 30546 | 30546 | 0.9999 |
| D1M7 | 51599 | 42042 | 33391 | 26808 | 26777 | 26777 | 0.9998 |
| D1M8 | 35464 | 28125 | 18372 | 15860 | 15667 | 15667 | 0.9997 |
| D1M9 | 46947 | 38306 | 28498 | 22621 | 22606 | 22606 | 0.9999 |
| D2M1 | 48895 | 38907 | 24908 | 17596 | 17197 | 17197 | 0.9999 |
| D2M10 | 37673 | 30395 | 24298 | 19097 | 19095 | 19095 | 0.9998 |
| D2M11 | 31545 | 25382 | 18525 | 16947 | 16834 | 16834 | 0.9999 |
| D2M12 | 65093 | 53174 | 43412 | 33439 | 33439 | 33439 | 0.9998 |
| D2M13 | 71186 | 58188 | 46114 | 33761 | 33761 | 33761 | 0.9999 |
| D2M14 | 36957 | 30161 | 24729 | 20702 | 20678 | 20702 | 1.0000 |
| D2M15 | 31876 | 25560 | 19673 | 19175 | 18498 | 18498 | 1.0000 |
| D2M16 | 43338 | 34911 | 27331 | 24121 | 24053 | 24053 | 1.0000 |
| D2M17 | 34741 | 28268 | 22337 | 17409 | 17409 | 17409 | 0.9999 |
| D2M18 | 26715 | 21305 | 16072 | 13539 | 13532 | 13532 | 1.0000 |
| D2M19 | 35641 | 28638 | 22589 | 19044 | 19044 | 19044 | 0.9997 |
| D2M2 | 30600 | 24300 | 17579 | 15002 | 14745 | 14745 | 1.0000 |
| D2M20 | 50963 | 40527 | 29316 | 24668 | 23979 | 23979 | 0.9999 |
| D2M21 | 44465 | 35977 | 28725 | 25127 | 25024 | 25024 | 0.9998 |
| D2M22 | 38953 | 31268 | 24501 | 20221 | 20215 | 20215 | 1.0000 |
| D2M23 | 20392 | 16228 | 13618 | 13526 | 12971 | 12979 | 0.9999 |
| D2M24 | 26952 | 21239 | 15275 | 12663 | 12539 | 12539 | 0.9999 |
| D2M25 | 43734 | 35497 | 25382 | 19984 | 19904 | 19904 | 0.9999 |
| D2M26 | 47088 | 37912 | 29000 | 22843 | 22843 | 22843 | 0.9999 |
| D2M27 | 36865 | 29752 | 23527 | 21456 | 21413 | 21440 | 1.0000 |
| D2M28 | 45776 | 36804 | 29730 | 28248 | 28057 | 28057 | 0.9999 |
| D2M29 | 33610 | 27325 | 21132 | 18304 | 18267 | 18267 | 1.0000 |
| D2M3 | 14446 | 11621 | 7613 | 5862 | 5845 | 5845 | 0.9997 |
| D2M30 | 32838 | 26143 | 17525 | 16293 | 15579 | 15579 | 0.9998 |
| D2M31 | 50633 | 40234 | 25995 | 23955 | 20862 | 20872 | 0.9996 |
| D2M32 | 25013 | 19807 | 16157 | 15959 | 15525 | 15525 | 0.9999 |
| D2M33 | 64812 | 52379 | 42308 | 41879 | 39529 | 39580 | 0.9999 |
| D2M34 | 50640 | 41106 | 33609 | 31132 | 31091 | 31111 | 0.9999 |
| D2M35 | 78468 | 64126 | 55413 | 52291 | 52162 | 52241 | 0.9999 |

| | | | | | | | |
|---|---|---|---|---|---|---|---|
| D2M36 | 39105 | 31818 | 22210 | 18752 | 18589 | 18589 | 0.9999 |
| D2M37 | 36270 | 29717 | 23941 | 19820 | 19820 | 19820 | 0.9997 |
| D2M38 | 50726 | 41452 | 33370 | 29995 | 29937 | 29945 | 0.9999 |
| D2M39 | 35836 | 29324 | 23131 | 17968 | 17968 | 17968 | 0.9999 |
| D2M4 | 23300 | 18897 | 15115 | 13036 | 13033 | 13033 | 0.9998 |
| D2M40 | 8197 | 6700 | 6029 | 5995 | 5529 | 5995 | 1.0000 |
| D2M41 | 42536 | 34706 | 25184 | 20067 | 20057 | 20057 | 1.0000 |
| D2M42 | 20766 | 16312 | 9379 | 8352 | 8201 | 8201 | 0.9998 |
| D2M43 | 48242 | 39105 | 28047 | 20988 | 20988 | 20988 | 0.9999 |
| D2M44 | 44157 | 35885 | 27168 | 19861 | 19861 | 19861 | 1.0000 |
| D2M45 | 30891 | 24870 | 16502 | 14084 | 13556 | 13556 | 0.9997 |
| D2M46 | 35631 | 29151 | 21937 | 16721 | 16721 | 16721 | 0.9999 |
| D2M47 | 42519 | 34912 | 28722 | 24448 | 24401 | 24430 | 0.9998 |
| D2M48 | 63931 | 51084 | 35450 | 28859 | 28351 | 28351 | 0.9999 |
| D2M5 | 22098 | 18050 | 13724 | 10894 | 10894 | 10894 | 0.9998 |
| D2M6 | 30431 | 24445 | 17364 | 12576 | 12572 | 12572 | 0.9998 |
| D2M7 | 39162 | 31598 | 23007 | 16923 | 16915 | 16915 | 0.9999 |
| D2M8 | 20861 | 16127 | 9223 | 7209 | 7103 | 7103 | 0.9999 |
| D2M9 | 45183 | 36852 | 27648 | 19978 | 19954 | 19954 | 0.9998 |
| D4M1 | 55653 | 43419 | 34490 | 29970 | 29631 | 29668 | 1.0000 |
| D4M10 | 33205 | 26221 | 22419 | 20254 | 20054 | 20122 | 1.0000 |
| D4M11 | 62770 | 49619 | 41368 | 40320 | 39873 | 40074 | 0.9999 |
| D4M12 | 37461 | 29351 | 25154 | 21810 | 21604 | 21656 | 0.9999 |
| D4M13 | 53905 | 41390 | 35882 | 25040 | 24990 | 25020 | 1.0000 |
| D4M14 | 33227 | 26164 | 21760 | 18657 | 18355 | 18419 | 0.9999 |
| D4M15 | 27089 | 21472 | 15033 | 13560 | 13006 | 13013 | 0.9999 |
| D4M16 | 66995 | 52244 | 40549 | 35021 | 34919 | 34925 | 1.0000 |
| D4M17 | 42850 | 33574 | 27619 | 19683 | 19665 | 19672 | 1.0000 |
| D4M18 | 47182 | 37597 | 30520 | 24366 | 24129 | 24129 | 1.0000 |
| D4M19 | 28887 | 22957 | 15813 | 13602 | 13516 | 13516 | 0.9999 |
| D4M2 | 36539 | 28459 | 22306 | 19316 | 19179 | 19184 | 0.9999 |
| D4M20 | 62384 | 49419 | 34662 | 25858 | 23273 | 23273 | 0.9999 |
| D4M21 | 50192 | 40140 | 31448 | 24784 | 24117 | 24117 | 0.9998 |
| D4M22 | 41268 | 33368 | 26290 | 18160 | 17838 | 17848 | 0.9999 |
| D4M23 | 50810 | 40249 | 27584 | 22905 | 22464 | 22466 | 0.9999 |
| D4M24 | 42695 | 33320 | 24926 | 19832 | 19684 | 19684 | 0.9998 |
| D4M25 | 49963 | 39513 | 30563 | 24680 | 23390 | 23390 | 0.9999 |
| D4M26 | 47639 | 36775 | 29551 | 17785 | 17701 | 17701 | 1.0000 |
| D4M27 | 49066 | 38708 | 29429 | 24746 | 24558 | 24558 | 1.0000 |
| D4M28 | 67115 | 53284 | 41277 | 38469 | 38105 | 38113 | 1.0000 |
| D4M29 | 56503 | 44567 | 35957 | 28361 | 27235 | 27235 | 1.0000 |
| D4M3 | 47320 | 36353 | 30448 | 23938 | 23143 | 23143 | 1.0000 |
| D4M30 | 31262 | 25149 | 15515 | 13415 | 9358 | 9358 | 0.9998 |

| | | | | | | | |
|---|---|---|---|---|---|---|---|
| D4M31 | 39989 | 31754 | 19843 | 17399 | 14482 | 14482 | 0.9999 |
| D4M32 | 38068 | 29461 | 21474 | 18789 | 18663 | 18667 | 0.9999 |
| D4M33 | 37740 | 28795 | 17076 | 14407 | 12874 | 12874 | 1.0000 |
| D4M34 | 31043 | 24407 | 18536 | 16164 | 16012 | 16012 | 1.0000 |
| D4M35 | 28615 | 21616 | 15202 | 12628 | 12480 | 12480 | 0.9998 |
| D4M36 | 39456 | 31263 | 25701 | 24108 | 23487 | 23487 | 0.9999 |
| D4M37 | 49670 | 37510 | 30863 | 20302 | 20293 | 20293 | 1.0000 |
| D4M38 | 42018 | 32441 | 25386 | 19530 | 19426 | 19426 | 0.9999 |
| D4M39 | 20328 | 15569 | 11925 | 8348 | 8329 | 8336 | 1.0000 |
| D4M4 | 41644 | 31674 | 26996 | 22070 | 21965 | 21974 | 1.0000 |
| D4M40 | 54919 | 42196 | 27049 | 22417 | 17626 | 17626 | 0.9999 |
| D4M41 | 54158 | 41564 | 32211 | 27007 | 26949 | 26949 | 0.9999 |
| D4M42 | 25541 | 19946 | 13757 | 11778 | 11271 | 11271 | 1.0000 |
| D4M43 | 50555 | 38705 | 31646 | 21141 | 21123 | 21123 | 1.0000 |
| D4M44 | 32879 | 25004 | 18390 | 13649 | 13624 | 13628 | 0.9999 |
| D4M45 | 42577 | 32780 | 20854 | 14501 | 8520 | 8520 | 0.9995 |
| D4M46 | 53392 | 42387 | 31063 | 26163 | 25968 | 25968 | 0.9999 |
| D4M47 | 33237 | 25990 | 19429 | 16014 | 15887 | 15887 | 0.9998 |
| D4M48 | 31424 | 24498 | 16640 | 12866 | 7941 | 7941 | 0.9997 |
| D4M5 | 47707 | 36866 | 31574 | 28523 | 28329 | 28350 | 1.0000 |
| D4M6 | 224290 | 174389 | 144420 | 119495 | 118515 | 118914 | 1.0000 |
| D4M7 | 76038 | 58829 | 49524 | 37355 | 37217 | 37223 | 1.0000 |
| D4M8 | 246510 | 189584 | 151548 | 125844 | 121963 | 122028 | 1.0000 |
| D4M9 | 27397 | 21656 | 18368 | 15844 | 15116 | 15116 | 1.0000 |
| D8MA | 287553 | 218524 | 180855 | 102424 | 99751 | 102336 | 1.0000 |
| D8M1 | 38435 | 30622 | 25789 | 22866 | 22668 | 22668 | 1.0000 |
| D8M10 | 29816 | 23620 | 18757 | 14818 | 14758 | 14758 | 1.0000 |
| D8M11 | 35079 | 26744 | 19552 | 15322 | 15244 | 15244 | 0.9999 |
| D8M12 | 49644 | 39332 | 35372 | 34263 | 33849 | 33869 | 1.0000 |
| D8M13 | 43759 | 34032 | 29592 | 25727 | 22679 | 22689 | 0.9998 |
| D8M14 | 42013 | 33416 | 28262 | 23585 | 23375 | 23375 | 1.0000 |
| D8M15 | 50396 | 39439 | 28233 | 26690 | 25990 | 25999 | 1.0000 |
| D8M16 | 43695 | 33867 | 27582 | 24958 | 24662 | 24674 | 1.0000 |
| D8M17 | 31532 | 24732 | 19827 | 15581 | 15545 | 15545 | 1.0000 |
| D8M18 | 39265 | 30791 | 24777 | 20845 | 20771 | 20771 | 1.0000 |
| D8M19 | 52814 | 40917 | 37614 | 37540 | 36862 | 37222 | 1.0000 |
| D8M2 | 35252 | 27744 | 20737 | 16723 | 16634 | 16634 | 0.9999 |
| D8M20 | 30477 | 24049 | 18960 | 16378 | 14878 | 14883 | 1.0000 |
| D8M21 | 47030 | 35062 | 28651 | 24125 | 23961 | 23968 | 1.0000 |
| D8M22 | 42448 | 34075 | 29516 | 28251 | 27650 | 28066 | 1.0000 |
| D8M23 | 39927 | 31194 | 21469 | 20216 | 19554 | 19652 | 0.9999 |
| D8M24 | 49526 | 38894 | 33760 | 33184 | 31941 | 32845 | 0.9999 |
| D8M25 | 33323 | 25918 | 21459 | 18116 | 17561 | 17561 | 1.0000 |

| | | | | | | | |
|---|---|---|---|---|---|---|---|
| D8M26 | 49710 | 39550 | 35151 | 34169 | 32861 | 33271 | 1.0000 |
| D8M27 | 61718 | 47906 | 38199 | 33446 | 33360 | 33360 | 1.0000 |
| D8M28 | 28683 | 22466 | 20225 | 20184 | 19388 | 19693 | 0.9999 |
| D8M29 | 46776 | 36845 | 32718 | 32164 | 31090 | 31926 | 1.0000 |
| D8M3 | 37076 | 29142 | 24166 | 21240 | 21036 | 21042 | 1.0000 |
| D8M30 | 41134 | 32398 | 26403 | 25870 | 22499 | 22992 | 1.0000 |
| D8M31 | 24944 | 19350 | 13878 | 13683 | 12482 | 12605 | 0.9999 |
| D8M32 | 40371 | 30824 | 22895 | 21481 | 21276 | 21407 | 1.0000 |
| D8M33 | 38454 | 29887 | 20529 | 18746 | 18226 | 18293 | 0.9999 |
| D8M34 | 54183 | 43616 | 35884 | 29525 | 29451 | 29473 | 1.0000 |
| D8M35 | 46849 | 37174 | 30666 | 26661 | 26278 | 26346 | 1.0000 |
| D8M36 | 31881 | 25528 | 23652 | 23595 | 21249 | 21871 | 1.0000 |
| D8M37 | 48265 | 38086 | 31704 | 26456 | 26124 | 26140 | 0.9999 |
| D8M38 | 71390 | 57861 | 50717 | 49386 | 48182 | 49060 | 1.0000 |
| D8M39 | 37241 | 30064 | 24839 | 20947 | 20737 | 20761 | 1.0000 |
| D8M4 | 50224 | 40439 | 37069 | 36908 | 36167 | 36759 | 0.9999 |
| D8M40 | 37598 | 29151 | 19300 | 15386 | 14210 | 14210 | 0.9999 |
| D8M41 | 40893 | 32143 | 25823 | 23333 | 23184 | 23215 | 1.0000 |
| D8M42 | 62225 | 49262 | 38975 | 34415 | 33444 | 33467 | 0.9999 |
| D8M43 | 44289 | 34670 | 28990 | 22919 | 22785 | 22785 | 1.0000 |
| D8M44 | 59895 | 46574 | 42987 | 42798 | 41805 | 42304 | 1.0000 |
| D8M45 | 38476 | 29691 | 22224 | 16308 | 15795 | 15810 | 1.0000 |
| D8M46 | 49975 | 39784 | 33307 | 28908 | 28850 | 28850 | 1.0000 |
| D8M47 | 66278 | 52652 | 46889 | 44139 | 43532 | 43578 | 0.9999 |
| D8M48 | 40737 | 31664 | 27462 | 25259 | 24153 | 24303 | 0.9999 |
| D8M5 | 45098 | 35950 | 31818 | 30245 | 29458 | 30032 | 1.0000 |
| D8M6 | 36420 | 29155 | 26892 | 26841 | 22829 | 25819 | 0.9999 |
| D8M7 | 50801 | 41370 | 36993 | 35691 | 34609 | 35359 | 1.0000 |
| D8M8 | 25787 | 19908 | 15227 | 13688 | 13466 | 13466 | 1.0000 |
| D8M9 | 23092 | 17849 | 13871 | 11813 | 11672 | 11672 | 1.0000 |
| D8MR | 234763 | 168907 | 137120 | 76100 | 71570 | 76003 | 0.9999 |
| D16MA | 211580 | 147602 | 113143 | 42070 | 35306 | 42067 | 0.9999 |
| D16M1 | 58724 | 41601 | 33768 | 28256 | 28198 | 28233 | 1.0000 |
| D16M10 | 57899 | 46167 | 38206 | 28545 | 27959 | 28358 | 0.9999 |
| D16M11 | 54169 | 42340 | 33573 | 27275 | 26905 | 26923 | 1.0000 |
| D16M12 | 73119 | 58863 | 49945 | 43799 | 43756 | 43775 | 1.0000 |
| D16M13 | 53374 | 42580 | 35426 | 25934 | 25795 | 25795 | 1.0000 |
| D16M14 | 52306 | 41877 | 35574 | 30382 | 30269 | 30357 | 1.0000 |
| D16M15 | 98739 | 74469 | 44107 | 34345 | 33329 | 33329 | 1.0000 |
| D16M16 | 30507 | 23316 | 17024 | 14520 | 14373 | 14373 | 1.0000 |
| D16M17 | 50702 | 40099 | 29995 | 21209 | 21110 | 21130 | 1.0000 |
| D16M18 | 47541 | 38220 | 29869 | 23757 | 23518 | 23537 | 0.9999 |
| D16M19 | 87 | 41 | 0 | 0 | 0 | 0 | 0.0000 |

| | | | | | | | |
|---|---|---|---|---|---|---|---|
| D16M2 | 27660 | 21073 | 14702 | 12646 | 12555 | 12569 | 1.0000 |
| D16M20 | 79694 | 56912 | 47265 | 37339 | 37334 | 37334 | 1.0000 |
| D16M21 | 57683 | 45969 | 38023 | 30151 | 30113 | 30113 | 1.0000 |
| D16M22 | 56743 | 45851 | 38913 | 32307 | 32305 | 32305 | 1.0000 |
| D16M23 | 60793 | 46423 | 27824 | 24750 | 23840 | 23848 | 1.0000 |
| D16M24 | 51788 | 40286 | 31828 | 28686 | 28425 | 28510 | 1.0000 |
| D16M25 | 61145 | 43327 | 34773 | 25998 | 25975 | 25975 | 1.0000 |
| D16M26 | 59371 | 46803 | 38052 | 31922 | 31901 | 31906 | 0.9999 |
| D16M27 | 52851 | 39315 | 21788 | 17709 | 17545 | 17545 | 0.9999 |
| D16M28 | 95554 | 72236 | 39132 | 32038 | 31691 | 31691 | 1.0000 |
| D16M29 | 51578 | 39312 | 31320 | 25573 | 25268 | 25274 | 1.0000 |
| D16M3 | 65307 | 50258 | 39821 | 31305 | 31104 | 31238 | 1.0000 |
| D16M30 | 43107 | 33222 | 24354 | 21602 | 20914 | 20946 | 1.0000 |
| D16M31 | 51736 | 40115 | 27114 | 24390 | 23809 | 23838 | 0.9999 |
| D16M32 | 67221 | 47647 | 25928 | 20695 | 20221 | 20221 | 1.0000 |
| D16M33 | 79661 | 61463 | 42338 | 35485 | 35099 | 35117 | 0.9999 |
| D16M34 | 66128 | 52250 | 37981 | 30382 | 30097 | 30097 | 1.0000 |
| D16M35 | 40382 | 31097 | 21644 | 17893 | 17711 | 17731 | 1.0000 |
| D16M36 | 71304 | 56904 | 47170 | 38108 | 38080 | 38080 | 1.0000 |
| D16M37 | 71629 | 57365 | 53226 | 53004 | 51874 | 52566 | 1.0000 |
| D16M38 | 53675 | 43198 | 35738 | 28637 | 28287 | 28287 | 0.9999 |
| D16M39 | 56131 | 44196 | 35825 | 27720 | 27636 | 27636 | 1.0000 |
| D16M4 | 90027 | 70944 | 59004 | 47171 | 47140 | 47140 | 1.0000 |
| D16M40 | 72982 | 57583 | 43963 | 38558 | 38017 | 38042 | 0.9999 |
| D16M41 | 60285 | 47082 | 37721 | 31629 | 31547 | 31614 | 1.0000 |
| D16M42 | 38853 | 31170 | 23458 | 18702 | 18558 | 18558 | 1.0000 |
| D16M43 | 48973 | 38510 | 33437 | 30426 | 30334 | 30378 | 1.0000 |
| D16M44 | 47411 | 37520 | 30794 | 23642 | 23512 | 23512 | 1.0000 |
| D16M45 | 57105 | 44024 | 33862 | 26582 | 26559 | 26559 | 1.0000 |
| D16M46 | 44027 | 34954 | 27582 | 24437 | 24141 | 24171 | 1.0000 |
| D16M47 | 45576 | 36071 | 28325 | 25472 | 25240 | 25295 | 1.0000 |
| D16M48 | 55601 | 43854 | 34643 | 27194 | 27021 | 27021 | 0.9999 |
| D16M5 | 76796 | 61202 | 50345 | 39982 | 39945 | 39945 | 1.0000 |
| D16M6 | 100242 | 79986 | 68793 | 57716 | 57557 | 57625 | 1.0000 |
| D16M7 | 49359 | 39433 | 32722 | 28822 | 28740 | 28783 | 1.0000 |
| D16M8 | 71152 | 55289 | 45571 | 40992 | 40876 | 40905 | 1.0000 |
| D16M9 | 77797 | 56894 | 47429 | 37294 | 37250 | 37256 | 1.0000 |
| D16MR | 104594 | 74869 | 58505 | 28093 | 16180 | 28079 | 1.0000 |
| D24MA | 201249 | 149068 | 124891 | 84573 | 72327 | 84571 | 1.0000 |
| D24M1 | 49711 | 40440 | 35476 | 33903 | 33611 | 33802 | 1.0000 |
| D24M10 | 68311 | 53971 | 44517 | 37523 | 29530 | 36426 | 1.0000 |
| D24M11 | 63324 | 50654 | 43309 | 39193 | 38495 | 38518 | 1.0000 |
| D24M12 | 53112 | 42202 | 35654 | 32493 | 32261 | 32352 | 1.0000 |

| | | | | | | | |
|---|---|---|---|---|---|---|---|
| D24M13 | 43456 | 34578 | 30840 | 29689 | 29538 | 29546 | 1.0000 |
| D24M14 | 45780 | 35972 | 30163 | 25307 | 25229 | 25229 | 1.0000 |
| D24M15 | 82255 | 63237 | 47524 | 46443 | 44588 | 44768 | 1.0000 |
| D24M16 | 79600 | 63617 | 52646 | 46370 | 45844 | 45854 | 1.0000 |
| D24M17 | 53626 | 43437 | 39234 | 38741 | 36186 | 36729 | 1.0000 |
| D24M18 | 54236 | 43591 | 35778 | 31375 | 31164 | 31213 | 1.0000 |
| D24M19 | 80772 | 64763 | 53606 | 51166 | 50585 | 50665 | 0.9999 |
| D24M2 | 2851 | 2279 | 2001 | 2001 | 1993 | 1993 | 1.0000 |
| D24M20 | 82764 | 65919 | 54822 | 45249 | 45203 | 45209 | 1.0000 |
| D24M21 | 39993 | 32345 | 27245 | 24538 | 24512 | 24512 | 1.0000 |
| D24M22 | 38341 | 31009 | 27872 | 27280 | 27185 | 27234 | 1.0000 |
| D24M23 | 69073 | 55509 | 43198 | 38131 | 37704 | 37852 | 0.9999 |
| D24M24 | 50533 | 38319 | 31909 | 29467 | 27215 | 27224 | 1.0000 |
| D24M25 | 99493 | 80220 | 67009 | 53168 | 53145 | 53148 | 1.0000 |
| D24M26 | 59692 | 47312 | 38495 | 30649 | 30383 | 30509 | 1.0000 |
| D24M27 | 70093 | 55449 | 41394 | 31976 | 31792 | 31792 | 1.0000 |
| D24M28 | 59437 | 46903 | 36604 | 30949 | 27246 | 27257 | 1.0000 |
| D24M29 | 69431 | 55361 | 44627 | 34169 | 34053 | 34053 | 1.0000 |
| D24M3 | 55096 | 44626 | 39679 | 38316 | 38153 | 38214 | 1.0000 |
| D24M30 | 88502 | 65994 | 53756 | 45356 | 42235 | 42238 | 0.9999 |
| D24M31 | 127112 | 98424 | 67588 | 54259 | 53496 | 53573 | 1.0000 |
| D24M32 | 125652 | 82618 | 65043 | 53731 | 50481 | 50693 | 0.9999 |
| D24M33 | 59043 | 46573 | 33787 | 26950 | 24116 | 24153 | 1.0000 |
| D24M34 | 33800 | 27017 | 21314 | 17535 | 16995 | 16995 | 1.0000 |
| D24M35 | 43026 | 34363 | 26907 | 21697 | 21346 | 21346 | 1.0000 |
| D24M36 | 68581 | 53253 | 43652 | 35975 | 35972 | 35972 | 1.0000 |
| D24M37 | 31814 | 25403 | 20160 | 15794 | 15715 | 15715 | 0.9999 |
| D24M38 | 51622 | 40183 | 34158 | 28586 | 28321 | 28321 | 1.0000 |
| D24M39 | 39382 | 31576 | 26007 | 21227 | 21046 | 21060 | 1.0000 |
| D24M4 | 51151 | 41436 | 36449 | 34539 | 34276 | 34296 | 1.0000 |
| D24M40 | 55660 | 43706 | 34917 | 32840 | 31572 | 31726 | 0.9999 |
| D24M41 | 69423 | 43457 | 33489 | 26762 | 25489 | 26678 | 1.0000 |
| D24M42 | 85531 | 64182 | 51319 | 45520 | 45346 | 45346 | 0.9999 |
| D24M43 | 74420 | 49561 | 37100 | 28961 | 28880 | 28880 | 1.0000 |
| D24M44 | 51834 | 40866 | 32815 | 26102 | 25594 | 25705 | 0.9999 |
| D24M45 | 48587 | 36893 | 28651 | 25885 | 25654 | 25698 | 0.9999 |
| D24M46 | 60415 | 47941 | 41611 | 36716 | 36053 | 36132 | 1.0000 |
| D24M47 | 67453 | 54489 | 44807 | 40459 | 39478 | 40129 | 1.0000 |
| D24M48 | 67978 | 54060 | 43394 | 37977 | 37697 | 37735 | 1.0000 |
| D24M5 | 41508 | 33204 | 27930 | 22304 | 22082 | 22082 | 1.0000 |
| D24M6 | 56793 | 45556 | 39907 | 37573 | 37181 | 37501 | 1.0000 |
| D24M7 | 76331 | 61525 | 51819 | 46427 | 46209 | 46289 | 1.0000 |
| D24M8 | 55399 | 43383 | 37493 | 35964 | 35634 | 35743 | 1.0000 |

| | | | | | | | |
|---|---|---|---|---|---|---|---|
| D24M9 | 53366 | 42939 | 38399 | 37538 | 37025 | 37406 | 1.0000 |
| D32MA | 185342 | 126284 | 99046 | 53636 | 50917 | 53616 | 1.0000 |
| D32M1 | 12339 | 5821 | 5026 | 4681 | 4668 | 4671 | 1.0000 |
| D32M10 | 34255 | 26079 | 19237 | 13131 | 10831 | 12806 | 0.9998 |
| D32M11 | 56965 | 40475 | 32406 | 26651 | 26548 | 26548 | 1.0000 |
| D32M12 | 35532 | 27206 | 21773 | 17509 | 16288 | 16300 | 1.0000 |
| D32M13 | 41607 | 32119 | 28109 | 24808 | 24768 | 24768 | 1.0000 |
| D32M14 | 36214 | 28301 | 25131 | 22820 | 22787 | 22791 | 1.0000 |
| D32M15 | 74244 | 56792 | 49308 | 47976 | 47336 | 47417 | 1.0000 |
| D32M16 | 22458 | 16064 | 13156 | 10569 | 10556 | 10556 | 1.0000 |
| D32M17 | 86954 | 66953 | 56691 | 54502 | 52820 | 53653 | 1.0000 |
| D32M18 | 66602 | 50945 | 42291 | 34726 | 34611 | 34647 | 1.0000 |
| D32M19 | 46320 | 35867 | 30252 | 25074 | 25046 | 25046 | 1.0000 |
| D32M2 | 92893 | 36740 | 29522 | 23602 | 23562 | 23584 | 1.0000 |
| D32M20 | 30777 | 15522 | 9470 | 7814 | 7273 | 7281 | 1.0000 |
| D32M21 | 47561 | 37114 | 32537 | 29760 | 28394 | 29760 | 1.0000 |
| D32M22 | 36473 | 28578 | 24280 | 21297 | 21149 | 21179 | 1.0000 |
| D32M23 | 38572 | 29701 | 21997 | 19159 | 19108 | 19108 | 0.9997 |
| D32M24 | 71288 | 45673 | 38894 | 35788 | 35695 | 35745 | 1.0000 |
| D32M25 | 86248 | 61270 | 51971 | 46888 | 46773 | 46869 | 1.0000 |
| D32M26 | 64013 | 49191 | 43123 | 39858 | 39653 | 39765 | 1.0000 |
| D32M27 | 53680 | 40859 | 33526 | 29757 | 29535 | 29535 | 1.0000 |
| D32M28 | 33571 | 24024 | 20084 | 18338 | 18233 | 18233 | 0.9999 |
| D32M29 | 21210 | 16519 | 14165 | 11767 | 11758 | 11758 | 1.0000 |
| D32M3 | 79891 | 63264 | 55154 | 48962 | 48820 | 48848 | 1.0000 |
| D32M30 | 39297 | 29166 | 21503 | 18233 | 17606 | 17606 | 0.9999 |
| D32M31 | 67000 | 51806 | 43914 | 42524 | 40294 | 41026 | 1.0000 |
| D32M32 | 42610 | 31554 | 27080 | 25705 | 25415 | 25577 | 1.0000 |
| D32M33 | 64333 | 49951 | 37146 | 27576 | 19428 | 19428 | 0.9999 |
| D32M34 | 88891 | 69507 | 60123 | 49333 | 49041 | 49041 | 1.0000 |
| D32M35 | 40409 | 31734 | 27519 | 25027 | 24804 | 24895 | 1.0000 |
| D32M36 | 25860 | 20352 | 16949 | 15886 | 15804 | 15860 | 1.0000 |
| D32M37 | 24802 | 18689 | 16176 | 16079 | 15888 | 15925 | 1.0000 |
| D32M38 | 33049 | 25372 | 21241 | 16312 | 16288 | 16288 | 1.0000 |
| D32M39 | 31975 | 25486 | 21897 | 19079 | 18844 | 18856 | 1.0000 |
| D32M4 | 91729 | 72954 | 63226 | 48293 | 48271 | 48271 | 1.0000 |
| D32M40 | 31330 | 23689 | 14995 | 12029 | 11993 | 11993 | 1.0000 |
| D32M41 | 55430 | 38130 | 30164 | 24205 | 23068 | 23486 | 1.0000 |
| D32M42 | 37703 | 25231 | 20428 | 17578 | 17512 | 17512 | 0.9999 |
| D32M43 | 36876 | 28198 | 23314 | 19777 | 18982 | 19037 | 1.0000 |
| D32M44 | 36696 | 26144 | 19989 | 16822 | 15684 | 15761 | 1.0000 |
| D32M45 | 29672 | 11987 | 6992 | 5702 | 5594 | 5597 | 1.0000 |
| D32M46 | 36888 | 29214 | 24652 | 21385 | 21033 | 21033 | 1.0000 |

| | | | | | | | |
|---|---|---|---|---|---|---|---|
| D32M47 | 35476 | 27893 | 23817 | 22226 | 21880 | 21916 | 1.0000 |
| D32M48 | 35750 | 24656 | 18242 | 15475 | 14723 | 14723 | 1.0000 |
| D32M5 | 30864 | 24760 | 20669 | 17485 | 17476 | 17476 | 1.0000 |
| D32M6 | 75250 | 59862 | 50371 | 43501 | 43434 | 43475 | 1.0000 |
| D32M7 | 50831 | 40080 | 34228 | 27488 | 27447 | 27447 | 1.0000 |
| D32M8 | 60451 | 37918 | 29067 | 22133 | 21828 | 21828 | 1.0000 |
| D32M9 | 35714 | 27587 | 23133 | 20949 | 20651 | 20903 | 1.0000 |
| D32MR | 51300 | 12482 | 9425 | 7111 | 6430 | 6812 | 1.0000 |
| D40MA | 194098 | 139322 | 117106 | 79639 | 60393 | 79591 | 1.0000 |
| D40M1 | 74155 | 56765 | 48217 | 42850 | 42695 | 42783 | 1.0000 |
| D40M10 | 79060 | 60008 | 52887 | 51212 | 48505 | 50614 | 1.0000 |
| D40M11 | 36732 | 26534 | 21973 | 17698 | 17667 | 17667 | 1.0000 |
| D40M12 | 24038 | 18402 | 14000 | 12139 | 10115 | 10207 | 1.0000 |
| D40M13 | 29225 | 23104 | 19166 | 14868 | 14830 | 14830 | 1.0000 |
| D40M14 | 56590 | 45148 | 39923 | 34864 | 34787 | 34790 | 1.0000 |
| D40M15 | 33994 | 26224 | 18252 | 15373 | 15337 | 15337 | 0.9999 |
| D40M16 | 51443 | 40017 | 32268 | 29451 | 29338 | 29338 | 1.0000 |
| D40M17 | 36896 | 28594 | 22422 | 17729 | 15626 | 15767 | 1.0000 |
| D40M18 | 58035 | 43577 | 35385 | 28061 | 27915 | 27991 | 0.9999 |
| D40M19 | 26220 | 20380 | 16939 | 15316 | 15264 | 15264 | 1.0000 |
| D40M2 | 44727 | 24228 | 18226 | 13649 | 13627 | 13644 | 1.0000 |
| D40M20 | 25257 | 18203 | 13167 | 10256 | 9990 | 9990 | 1.0000 |
| D40M21 | 18325 | 13699 | 10378 | 8785 | 8656 | 8785 | 1.0000 |
| D40M22 | 18865 | 14565 | 11778 | 10334 | 10258 | 10283 | 1.0000 |
| D40M23 | 18546 | 14195 | 9713 | 8883 | 8729 | 8736 | 1.0000 |
| D40M24 | 26590 | 20219 | 17405 | 15594 | 15590 | 15590 | 1.0000 |
| D40M25 | 31953 | 21195 | 12477 | 9467 | 9248 | 9248 | 1.0000 |
| D40M26 | 46736 | 35899 | 28378 | 25864 | 25725 | 25826 | 1.0000 |
| D40M27 | 34272 | 26306 | 19888 | 18850 | 18784 | 18784 | 0.9999 |
| D40M28 | 26700 | 19262 | 12755 | 11833 | 11782 | 11782 | 1.0000 |
| D40M29 | 27822 | 21817 | 15731 | 14600 | 14535 | 14545 | 0.9999 |
| D40M3 | 39225 | 31083 | 27567 | 22742 | 22732 | 22732 | 1.0000 |
| D40M30 | 25446 | 19101 | 10665 | 9619 | 9238 | 9238 | 0.9999 |
| D40M31 | 27695 | 21245 | 12525 | 11616 | 11176 | 11176 | 1.0000 |
| D40M32 | 18535 | 13941 | 10796 | 9559 | 9550 | 9550 | 1.0000 |
| D40M33 | 86634 | 68229 | 49983 | 41330 | 36816 | 36891 | 1.0000 |
| D40M34 | 63811 | 50363 | 42970 | 33582 | 33426 | 33426 | 0.9999 |
| D40M35 | 41489 | 33064 | 27873 | 24615 | 24546 | 24546 | 1.0000 |
| D40M36 | 27657 | 20337 | 15681 | 12217 | 11722 | 11722 | 0.9999 |
| D40M37 | 26443 | 21033 | 15922 | 12740 | 12098 | 12098 | 1.0000 |
| D40M38 | 38328 | 30866 | 27598 | 22965 | 22759 | 22890 | 1.0000 |
| D40M39 | 29396 | 22580 | 18117 | 15166 | 13967 | 13967 | 1.0000 |
| D40M4 | 27218 | 21350 | 18273 | 13913 | 13883 | 13897 | 0.9999 |

| | | | | | | | |
|---|---|---|---|---|---|---|---|
| D40M40 | 27850 | 20352 | 13750 | 12485 | 12421 | 12421 | 1.0000 |
| D40M41 | 35971 | 25691 | 22109 | 19841 | 19648 | 19831 | 1.0000 |
| D40M42 | 30996 | 18849 | 13820 | 10466 | 9230 | 9230 | 0.9999 |
| D40M43 | 17804 | 13476 | 10123 | 7668 | 7545 | 7545 | 1.0000 |
| D40M44 | 36415 | 27362 | 22542 | 20042 | 14667 | 14821 | 0.9999 |
| D40M45 | 168884 | 90157 | 73745 | 49821 | 49138 | 49138 | 1.0000 |
| D40M46 | 19232 | 14813 | 12427 | 11665 | 11511 | 11511 | 1.0000 |
| D40M47 | 24782 | 18741 | 15070 | 13796 | 13624 | 13624 | 1.0000 |
| D40M48 | 23189 | 17423 | 12541 | 11761 | 10963 | 10963 | 1.0000 |
| D40M5 | 23726 | 18312 | 15509 | 12745 | 12732 | 12745 | 1.0000 |
| D40M6 | 37987 | 29221 | 24218 | 19274 | 19261 | 19274 | 1.0000 |
| D40M7 | 38304 | 30345 | 25256 | 17930 | 17858 | 17892 | 1.0000 |
| D40M8 | 32481 | 20646 | 14897 | 11982 | 11962 | 11962 | 1.0000 |
| D40M9 | 88578 | 70746 | 60251 | 45263 | 45032 | 45245 | 1.0000 |
| D40MR | 141960 | 67090 | 40888 | 15902 | 12506 | 15875 | 0.9997 |
| D48MA | 228608 | 69182 | 31238 | 12703 | 11197 | 12703 | 0.9994 |
| D48M1 | 31329 | 23269 | 21221 | 21204 | 20511 | 21033 | 1.0000 |
| D48M10 | 44871 | 34696 | 25717 | 19730 | 18257 | 19150 | 1.0000 |
| D48M11 | 50186 | 36129 | 29180 | 24416 | 24372 | 24372 | 1.0000 |
| D48M12 | 37297 | 28637 | 22184 | 18866 | 16714 | 17035 | 1.0000 |
| D48M13 | 73690 | 55855 | 47701 | 39188 | 39042 | 39042 | 1.0000 |
| D48M14 | 49742 | 39304 | 33061 | 27159 | 27051 | 27088 | 0.9999 |
| D48M15 | 25769 | 20035 | 17951 | 17852 | 17056 | 17786 | 0.9999 |
| D48M16 | 62113 | 48848 | 41395 | 36283 | 36249 | 36266 | 1.0000 |
| D48M17 | 31393 | 24652 | 19287 | 14859 | 13264 | 13297 | 1.0000 |
| D48M18 | 43105 | 34580 | 29887 | 28545 | 28317 | 28327 | 1.0000 |
| D48M19 | 28024 | 21959 | 16762 | 12166 | 12115 | 12152 | 1.0000 |
| D48M2 | 32940 | 22841 | 18719 | 17525 | 17459 | 17508 | 1.0000 |
| D48M20 | 31202 | 24280 | 15717 | 11710 | 11329 | 11329 | 0.9999 |
| D48M21 | 35908 | 27929 | 20965 | 16078 | 16007 | 16072 | 1.0000 |
| D48M22 | 44555 | 34795 | 26676 | 17237 | 16506 | 16517 | 1.0000 |
| D48M23 | 52256 | 41448 | 31457 | 26469 | 26069 | 26085 | 1.0000 |
| D48M24 | 32007 | 21349 | 17641 | 15051 | 15051 | 15051 | 1.0000 |
| D48M25 | 46364 | 33139 | 27697 | 20753 | 20736 | 20742 | 1.0000 |
| D48M26 | 33747 | 26504 | 21824 | 18733 | 18531 | 18600 | 0.9999 |
| D48M27 | 56360 | 43344 | 36047 | 29707 | 29682 | 29682 | 1.0000 |
| D48M28 | 37840 | 26094 | 20694 | 17863 | 17721 | 17737 | 0.9999 |
| D48M29 | 1560 | 1118 | 780 | 757 | 752 | 752 | 1.0000 |
| D48M3 | 34409 | 26735 | 23705 | 21435 | 21416 | 21416 | 1.0000 |
| D48M30 | 28976 | 22382 | 14837 | 12635 | 11084 | 11084 | 1.0000 |
| D48M31 | 20971 | 16465 | 12204 | 10541 | 10235 | 10243 | 1.0000 |
| D48M32 | 50265 | 30119 | 22739 | 18326 | 18250 | 18280 | 1.0000 |
| D48M33 | 31334 | 23217 | 15500 | 11596 | 10562 | 10562 | 0.9999 |

| | | | | | | | |
|---|---|---|---|---|---|---|---|
| D48M34 | 33383 | 26203 | 21212 | 17338 | 17205 | 17205 | 1.0000 |
| D48M35 | 34411 | 26273 | 21483 | 17730 | 17659 | 17659 | 1.0000 |
| D48M36 | 24400 | 17786 | 13852 | 10040 | 8963 | 8963 | 1.0000 |
| D48M37 | 36075 | 27226 | 20358 | 15303 | 14879 | 14879 | 1.0000 |
| D48M38 | 40011 | 30278 | 24577 | 18227 | 17742 | 18005 | 1.0000 |
| D48M39 | 33031 | 24918 | 19339 | 14140 | 12623 | 12623 | 1.0000 |
| D48M4 | 31071 | 24432 | 21952 | 19578 | 19496 | 19496 | 1.0000 |
| D48M40 | 46094 | 33824 | 19579 | 15131 | 14768 | 14793 | 0.9998 |
| D48M41 | 45430 | 30663 | 25246 | 21232 | 21024 | 21201 | 1.0000 |
| D48M42 | 38818 | 29085 | 17783 | 13311 | 13074 | 13074 | 0.9998 |
| D48M43 | 22949 | 17435 | 13920 | 10748 | 10166 | 10166 | 1.0000 |
| D48M44 | 44408 | 33298 | 26822 | 21855 | 18171 | 18605 | 1.0000 |
| D48M45 | 37594 | 27233 | 21406 | 16447 | 16251 | 16251 | 1.0000 |
| D48M46 | 28910 | 22613 | 18057 | 15238 | 14843 | 14843 | 0.9999 |
| D48M47 | 29140 | 22584 | 17679 | 14553 | 14188 | 14188 | 1.0000 |
| D48M48 | 14644 | 10819 | 7359 | 6242 | 5021 | 5034 | 1.0000 |
| D48M5 | 49442 | 38543 | 33521 | 30610 | 30610 | 30610 | 1.0000 |
| D48M6 | 56929 | 45144 | 39354 | 36430 | 36423 | 36423 | 1.0000 |
| D48M7 | 60041 | 47279 | 42015 | 39128 | 38339 | 38641 | 1.0000 |
| D48M8 | 65408 | 40130 | 31857 | 24866 | 24797 | 24797 | 1.0000 |
| D48M9 | 40055 | 30864 | 24868 | 17926 | 17828 | 17852 | 1.0000 |
| D48MR | 240704 | 85759 | 34713 | 15050 | 13263 | 15050 | 0.9995 |
| D56MA | 31399 | 10625 | 5788 | 4178 | 2922 | 4178 | 1.0000 |
| D56M1 | 47980 | 37076 | 31185 | 27117 | 26862 | 26964 | 1.0000 |
| D56M10 | 50203 | 37824 | 30592 | 22983 | 21759 | 22408 | 0.9999 |
| D56M11 | 39632 | 26887 | 21542 | 19268 | 19233 | 19247 | 1.0000 |
| D56M12 | 31620 | 23728 | 18809 | 15891 | 13740 | 14188 | 1.0000 |
| D56M13 | 105 | 50 | 5 | 5 | 5 | 5 | 1.0000 |
| D56M14 | 101394 | 77416 | 66796 | 51578 | 48747 | 49951 | 0.9999 |
| D56M15 | 25179 | 18626 | 11429 | 9882 | 9858 | 9873 | 1.0000 |
| D56M16 | 33411 | 25618 | 20806 | 18132 | 18132 | 18132 | 1.0000 |
| D56M17 | 41663 | 31812 | 28147 | 28127 | 26971 | 27489 | 1.0000 |
| D56M18 | 38350 | 29842 | 24414 | 22607 | 22503 | 22524 | 1.0000 |
| D56M19 | 38510 | 28668 | 23359 | 20528 | 20003 | 20521 | 1.0000 |
| D56M2 | 91478 | 71750 | 59644 | 47611 | 47442 | 47611 | 1.0000 |
| D56M20 | 27622 | 20653 | 15144 | 12861 | 12837 | 12837 | 1.0000 |
| D56M21 | 25042 | 18795 | 15025 | 11841 | 11825 | 11841 | 1.0000 |
| D56M22 | 40764 | 31960 | 26714 | 20664 | 20395 | 20428 | 1.0000 |
| D56M23 | 27282 | 21060 | 14670 | 12953 | 12749 | 12749 | 0.9995 |
| D56M24 | 74367 | 57385 | 49224 | 44157 | 44157 | 44157 | 1.0000 |
| D56M25 | 45932 | 27750 | 17271 | 14476 | 14463 | 14463 | 1.0000 |
| D56M26 | 20881 | 16380 | 14468 | 13820 | 13661 | 13685 | 1.0000 |
| D56M27 | 69002 | 52379 | 45542 | 42222 | 42192 | 42204 | 1.0000 |

| | | | | | | | |
|---|---|---|---|---|---|---|---|
| D56M28 | 75788 | 57615 | 47487 | 42257 | 41044 | 41044 | 1.0000 |
| D56M29 | 30285 | 23148 | 18718 | 16410 | 16349 | 16349 | 1.0000 |
| D56M3 | 57143 | 43292 | 37180 | 31140 | 31061 | 31061 | 1.0000 |
| D56M30 | 48222 | 37128 | 28037 | 22377 | 21155 | 21159 | 1.0000 |
| D56M31 | 45736 | 34919 | 24613 | 21030 | 20385 | 20385 | 1.0000 |
| D56M32 | 21085 | 15821 | 13017 | 11607 | 11564 | 11564 | 1.0000 |
| D56M33 | 75596 | 58518 | 44239 | 34386 | 29479 | 29516 | 1.0000 |
| D56M34 | 61830 | 49072 | 40581 | 32628 | 32376 | 32376 | 1.0000 |
| D56M35 | 39970 | 31162 | 26037 | 23852 | 23780 | 23812 | 1.0000 |
| D56M36 | 37657 | 29304 | 24753 | 19650 | 18649 | 18649 | 0.9999 |
| D56M37 | 63621 | 50229 | 42775 | 33036 | 32164 | 32169 | 1.0000 |
| D56M38 | 73994 | 58081 | 50646 | 46488 | 45101 | 46011 | 1.0000 |
| D56M39 | 62717 | 49813 | 41901 | 32634 | 30701 | 30701 | 1.0000 |
| D56M4 | 29728 | 23295 | 20901 | 19698 | 19448 | 19456 | 1.0000 |
| D56M40 | 48568 | 36095 | 27561 | 21777 | 19802 | 19802 | 0.9999 |
| D56M41 | 119092 | 92072 | 78197 | 65571 | 65092 | 65493 | 0.9999 |
| D56M42 | 26534 | 20665 | 16333 | 14470 | 10478 | 10714 | 1.0000 |
| D56M43 | 60909 | 47006 | 40758 | 33184 | 32540 | 32568 | 1.0000 |
| D56M44 | 35188 | 27340 | 23810 | 20231 | 18375 | 18755 | 1.0000 |
| D56M45 | 37168 | 26629 | 21457 | 17299 | 17182 | 17201 | 1.0000 |
| D56M46 | 47600 | 37987 | 33790 | 30666 | 30586 | 30586 | 1.0000 |
| D56M47 | 71163 | 56737 | 48719 | 42688 | 42571 | 42587 | 1.0000 |
| D56M48 | 25149 | 18743 | 18002 | 18002 | 16845 | 18002 | 1.0000 |
| D56M5 | 39186 | 30426 | 26580 | 25104 | 24503 | 24511 | 1.0000 |
| D56M6 | 76622 | 59258 | 48645 | 38826 | 38193 | 38228 | 0.9999 |
| D56M7 | 71254 | 55730 | 48263 | 39548 | 37543 | 37595 | 1.0000 |
| D56M8 | 25865 | 15557 | 12543 | 11149 | 11084 | 11084 | 1.0000 |
| D56M9 | 35189 | 25910 | 21591 | 17721 | 17291 | 17373 | 1.0000 |
| D56MR | 149917 | 98082 | 77874 | 56515 | 40421 | 56366 | 0.9999 |
| D64M1 | 29854 | 14245 | 6723 | 5849 | 5849 | 5849 | 1.0000 |
| D64M10 | 44410 | 21620 | 8029 | 6395 | 6136 | 6395 | 0.9997 |
| D64M11 | 34773 | 14829 | 6120 | 5012 | 5005 | 5005 | 1.0000 |
| D64M12 | 39109 | 16271 | 4285 | 2969 | 2893 | 2969 | 0.9997 |
| D64M13 | 62298 | 28067 | 9555 | 8044 | 5067 | 8044 | 1.0000 |
| D64M14 | 46096 | 22057 | 6979 | 5057 | 4242 | 5057 | 1.0000 |
| D64M15 | 38530 | 16606 | 4634 | 4181 | 4181 | 4181 | 1.0000 |
| D64M16 | 35933 | 16423 | 5163 | 4785 | 4770 | 4770 | 1.0000 |
| D64M17 | 35911 | 17034 | 4081 | 3354 | 3351 | 3354 | 1.0000 |
| D64M18 | 65693 | 32382 | 15188 | 13944 | 13916 | 13916 | 1.0000 |
| D64M19 | 60574 | 28727 | 14193 | 12730 | 12367 | 12730 | 1.0000 |
| D64M2 | 51600 | 25812 | 11581 | 9195 | 9166 | 9166 | 0.9998 |
| D64M20 | 51793 | 24818 | 7483 | 7008 | 7004 | 7004 | 1.0000 |
| D64M21 | 59300 | 31807 | 18280 | 12919 | 12919 | 12919 | 0.9998 |

| | | | | | | | | |
|---|---|---|---|---|---|---|---|---|
| D64M22 | 42513 | 21258 | 9655 | 8350 | | 8350 | 8350 | 1.0000 |
| D64M23 | 39495 | 17961 | 6031 | 5490 | | 5355 | 5364 | 1.0000 |
| D64M24 | 45800 | 22936 | 12107 | 11453 | | 11453 | 11453 | 1.0000 |
| D64M25 | 40281 | 19541 | 8885 | 7964 | | 7964 | 7964 | 1.0000 |
| D64M26 | 53127 | 26833 | 13260 | 12344 | | 12344 | 12344 | 1.0000 |
| D64M27 | 34869 | 16514 | 5262 | 4480 | | 4480 | 4480 | 1.0000 |
| D64M28 | 33478 | 15667 | 6713 | 5832 | | 5729 | 5729 | 1.0000 |
| D64M29 | 53034 | 27358 | 13692 | 12759 | | 12759 | 12759 | 1.0000 |
| D64M3 | 36841 | 18383 | 8506 | 7461 | | 7461 | 7461 | 1.0000 |
| D64M30 | 47916 | 22295 | 8853 | 6941 | | 6882 | 6882 | 1.0000 |
| D64M31 | 40007 | 20851 | 2982 | 2580 | | 2546 | 2546 | 1.0000 |
| D64M32 | 38862 | 17836 | 9364 | 8449 | | 8402 | 8402 | 1.0000 |
| D64M33 | 37430 | 20071 | 2574 | 2214 | | 2067 | 2067 | 1.0000 |
| D64M34 | 52008 | 25490 | 11077 | 9511 | | 9448 | 9511 | 0.9999 |
| D64M35 | 34042 | 16692 | 6002 | 4982 | | 4982 | 4982 | 1.0000 |
| D64M36 | 19544 | 9869 | 4205 | 3900 | | 3892 | 3900 | 1.0000 |
| D64M37 | 31539 | 15470 | 3843 | 3064 | | 3064 | 3064 | 1.0000 |
| D64M38 | 21365 | 10103 | 6034 | 5851 | | 5851 | 5851 | 1.0000 |
| D64M39 | 25309 | 12264 | 4225 | 3189 | | 3189 | 3189 | 1.0000 |
| D64M4 | 48658 | 23032 | 7271 | 6295 | | 6290 | 6295 | 1.0000 |
| D64M40 | 29948 | 13296 | 5648 | 4147 | | 4126 | 4126 | 0.9998 |
| D64M41 | 39199 | 17878 | 9438 | 7557 | | 7557 | 7557 | 1.0000 |
| D64M42 | 37176 | 18005 | 7837 | 6798 | | 5759 | 5759 | 1.0000 |
| D64M43 | 38559 | 19067 | 6141 | 4673 | | 4673 | 4673 | 0.9998 |
| D64M44 | 37969 | 17331 | 4417 | 3543 | | 3453 | 3543 | 1.0000 |
| D64M45 | 44059 | 18632 | 2829 | 2527 | | 2527 | 2527 | 1.0000 |
| D64M46 | 22816 | 10960 | 6330 | 6019 | | 6019 | 6019 | 1.0000 |
| D64M47 | 26436 | 12192 | 5003 | 4634 | | 4626 | 4626 | 1.0000 |
| D64M48 | 35172 | 13658 | 2568 | 2342 | | 2276 | 2276 | 1.0000 |
| D64M5 | 44378 | 20787 | 4743 | 3940 | | 3783 | 3940 | 1.0000 |
| D64M6 | 36044 | 16190 | 3590 | 2793 | | 2759 | 2793 | 1.0000 |
| D64M7 | 48272 | 23916 | 8232 | 7067 | | 7060 | 7067 | 1.0000 |
| D64M8 | 58491 | 25723 | 12363 | 11542 | | 11542 | 11542 | 0.9999 |
| D64M9 | 42847 | 21426 | 11020 | 7083 | | 6929 | 7083 | 1.0000 |
| D64MAir | 38199 | 18514 | 6740 | 4426 | | 2916 | 4426 | 1.0000 |
| D64MRain | 26851 | 13791 | 5045 | 3796 | | 2664 | 3796 | 1.0000 |
| D64MextractionMnegM1 | 154 | 32 | 0 | 0 | NA | | 0 | |
| D64MextractionMnegM2 | 181 | 33 | 0 | 0 | NA | | 0 | |
| D64MPCRMneg | 118 | 27 | 0 | 0 | NA | | 0 | |
| D64MPCRMneg2 | 85 | 11 | 0 | 0 | NA | | 0 | |
| ExtractionMneg1P1 | 3876 | 2692 | 2269 | 2266 | NA | | 2228 | |
| ExtractionMneg1P2 | 316 | 189 | 86 | 86 | NA | | 86 | |
| ExtractionMneg2P1 | 3410 | 2282 | 1626 | 1626 | NA | | 1570 | |

| | | | | | | |
|---|---|---|---|---|---|---|
| ExtractionMneg2P2 | 54 | 28 | 0 | 0 | NA | 0 |
| PCRMneg1P1 | 17294 | 13122 | 9034 | 6712 | NA | 6410 |
| PCRMneg1P2 | 5561 | 3878 | 2087 | 1667 | NA | 1606 |
| PCRMneg2P1 | 61 | 38 | 21 | 21 | NA | 21 |
| PCRMneg2P2 | 246 | 145 | 38 | 38 | NA | 38 |
